# Conserved molecular recognition by an intrinsically disordered region in the absence of sequence conservation

**DOI:** 10.1101/2023.08.06.552128

**Authors:** Jhullian J. Alston, Andrea Soranno, Alex S. Holehouse

## Abstract

Intrinsically disordered regions (IDRs) are critical for cellular function, yet often appear to lack sequence conservation when assessed by multiple sequence alignments. This raises the question of if and how function can be encoded and preserved in these regions despite massive sequence variation. To address this question, we have applied coarse-grained molecular dynamics simulations to investigate non-specific RNA binding of coronavirus nucleocapsid proteins. Coronavirus nucleocapsid proteins consist of multiple interspersed disordered and folded domains that bind RNA. We focussed here on the first two domains of coronavirus nucleocapsid proteins, the disordered N-terminal domain (NTD) followed by the folded RNA binding domain (RBD). While the NTD is highly variable across evolution, the RBD is structurally conserved. This combination makes the NTD-RBD a convenient model system to explore the interplay between an IDR adjacent to a folded domain, and how changes in IDR sequence can influence molecular recognition of a partner. Our results reveal a surprising degree of sequence-specificity encoded by both the composition and the precise order of the amino acids in the NTD. The presence of an NTD can – depending on the sequence – either suppress or enhance RNA binding. Despite this sensitivity, large-scale variation in NTD sequences is possible while certain sequence features are retained. Consequently, a conformationally-conserved fuzzy RNA:protein complex is found across nucleocapsid protein orthologs, despite large-scale changes in both NTD sequence and RBD surface chemistry. Taken together, these insights shed light on the ability of disordered regions to preserve functional characteristics despite their sequence variability.

## Introduction

The classical structure-function paradigm states that sequence dictates structure, and structure dictates function(1). This understanding has driven extensive study of protein structure and dynamics. Understanding the 3D structures that proteins adopt provides insight into their normal function. It also allows us to interpret how and why mutations that disrupt those structures and/or dynamics impair function(2–4). However, in recent years, there has been a growing focus on understanding “unstructured” or disordered protein regions(5–8). Intrinsically disordered regions (IDRs) are poorly described by a single 3D structure; instead, they exist as a collection of structurally distinct interconverting conformations known as an ensemble (9, 10). Despite lacking a defined 3D structure, IDRs play critical roles in many aspects of cellular function. Consequently, emerging work suggests that just as folded domains follow a sequence-structure-function relationship, IDRs can follow an analogous sequence-ensemble-function relationship (11). Given the importance that structure-function analysis has played in understanding the molecular basis for cellular function, there is a promising and analogous opportunity to understand IDR function through the lens of ensembles (12–16).

A major goal of modern molecular biology is to accurately predict protein function directly from amino acid sequence. Rooted in the general assumption that similar protein sequences will exhibit similar molecular behavior, one strategy is to compare the sequence of a protein of interest to those of other known proteins (17–20). In many cases, multiple sequence alignment of orthologous folded domains reveals high sequence conservation and, therefore, conserved protein function. (17, 21, 22). This relationship enables us to predict structures of previously unsolved protein structures and infer function by aligning the sequences of an uncharacterized protein against sequences of functionally-characterized folded domains (23–25). In sum, applying evolutionary information, directly and indirectly, is a central pillar in our modern toolkit for protein sequence analysis.

While IDR sequences can be aligned, their conservation at the residue level is typically lower than their structured counterparts(26–28). However, even without strict sequence conservation, the presence of disorder in a given protein domain is often conserved across orthologs(12, 13, 28–31). Assuming orthologous proteins provide equivalent functions, this presents an intriguing question: “Can apparently divergent IDRs confer the same molecular functions?””. For some IDRs, the only feature that matters may be the existence of Short Linear Motifs (SLiMs), such that a large IDR may appear poorly conserved, yet functional conservation is maintained as long as a few short (5-15 residue) regions are present(32–34). Recent work suggests that retaining specific physicochemical properties in a disordered region is sufficient to preserve function(13, 26, 28–30, 35–37). Some of the conformations disordered proteins may adopt can be structured. This transient structure formation can underlie conservation in some IDRs, where specific interactions are needed to maintain proper folds for protein-protein and protein-nucleic acid interactions(14, 38). Ultimately, the absence of a specific 3D structure loosens the relationship between sequence and function.

Viruses provide good test systems for exploring evolutionary conservation in IDRs. Viruses use IDRs extensively, and their rapid evolutionary rates – driven by a combination of fast replication times, massive numbers, and strong fitness selection – mean that even between serotypes of the same virus, substantial divergence in IDRs is often observed (39–44). For viruses that infect the same host, it is reasonable to expect equivalent selective pressures and equivalent protein function. Taken together, viral IDRs offer a convenient opportunity to explore how large-scale variation in IDR sequence enables similar functional output.

In this work, we investigated the relationship between IDR sequence and RNA interaction by performing coarse-grained molecular dynamics simulations of coronavirus nucleocapsid (N) proteins(45, 46). Coronaviruses are positive-sense single-stranded RNA viruses with relatively large (∼30 Kb) genomes (46–49). They typically consist of 4 major structural proteins: spike (S), envelope (E), membrane (M), and the N protein. The N protein is the most abundant viral protein and drives genomic RNA condensation and packaging during virion assembly, but has also been implicated in the evasion of the host immune system(50–53). Given its abundance and importance, the N protein is a tractable model system for exploring variation in sequence and function.

Coronavirus N proteins consist of five domains; two folded domains and three IDRs (**Fig. 1A**) (54). Our prior work systematically characterized full-length SARS-CoV-2 N protein using a combination of all-atom simulations, single-molecule Förster Resonance Energy Transfer (smFRET) spectroscopy, and nanosecond Fluorescence Correlation Spectroscopy (ns-FCS) (51). This work confirmed the disordered nature of the three IDRs and characterized their ensemble behavior in the context of the full-length protein. The two N-terminal domains (the N-terminal domain, NTD, and RNA-binding domain, RBD) are disordered and folded, respectively. In addition to characterization in the absence of RNA, our more recent experimental and computational work showed that these domains work together to enable high-affinity RNA binding (52). While the RBD alone binds (rU)_25_ with a binding affinity of ∼0.6 µM^-1^, the addition of the NTD enhances this affinity around 30-fold. This enhancement in binding affinity is facilitated by a fuzzy complex that forms between the NTD-RBD and RNA, where the NTD remains fully disordered in the bound and unbound states. While we cannot exclude other potential roles for the NTD, our work to date suggests that one of its functions is to enhance N-protein:RNA interactions, presumably to facilitate genome packaging.

**Figure 1.**
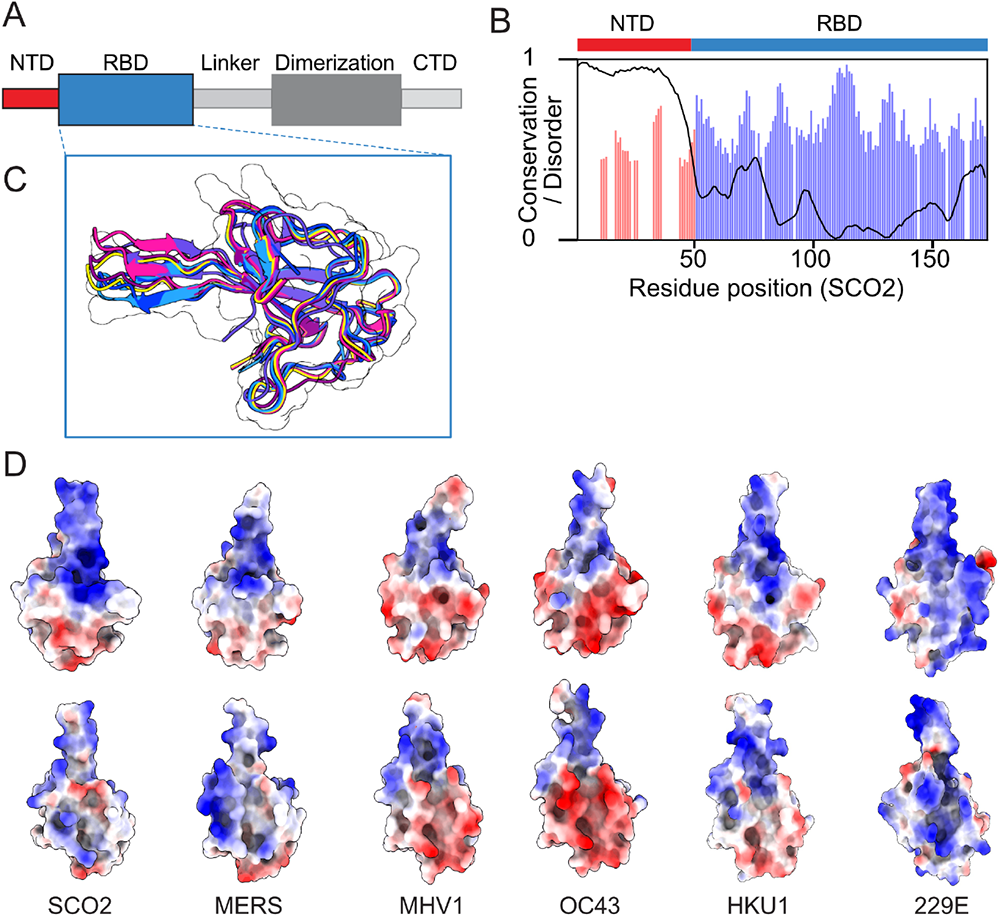
Coronavirus nucleocapsid proteins possess a disordered, poorly-conserved N-terminal domain (NTD) and a more well-conserved folded RNA binding domain (RBD). **A.** Schematic showing full-length nucleocapsid protein architectures from coronaviruses. The nucleocapsid protein contains three IDRs (NTD, Linker, CTD) and two folded domains (RBD, and Dimerization domains). **B.** Per-residue conservation calculated over 45 orthologous NTR-RBD constructs, including SCO2, MERS, OC43, HKU1, 229E, and MHV1. Conservation is calculated based on the positional Shannon entropy, with values shown only for residues where 80% or more of orthologous possess a residue. The NTD contains many gaps in a relatively poor alignment, while the RBD is almost uniformly populated with relatively highly conserved residues. **C.** Overlay of RBD structures for SCO2, MERS, OC43, HKU1, 229E, and MHV1, revealing a high degree of structural conservation in the RBD fold. **D.** Surface charge properties of the six RBD structures overlaid in panel C, highlighting differences in surface charge properties despite the conservation of the overall fold.

In this work, we focussed on the NTD and RBD as a model system for understanding the sequence constraints on molecular function. While the NTD sequence is variable across N protein orthologs, the presence of a disordered NTD is highly conserved in coronaviruses (**Fig. 1B**) (51). In contrast, the RBD is highly conserved among orthologs, and, despite some variation in sequence leading to changes in surface chemistry, it harbors a nearly identical fold across experimentally resolved and computationally predicted structures (**Fig. 1C**).

Given the structurally similar RBDs but differing NTDs, we wondered whether different coronavirus NTD-RBDs bind single-stranded RNA (ssRNA) in the same way, or whether they have distinct modes of interaction. Naively, given the large variation in NTD sequence, one might expect fundamentally different modes of recognition. However, recent work has shown that the conservation of IDR ensemble properties is possible despite large changes in IDR sequence(12).

To address this question, we performed coarse-grained molecular dynamics (MD) simulations of NTD-RBD constructs with poly-(rU)_25_ to assess how changes in NTD sequence influence molecular function, i.e., RNA binding. Using this approach, we sought to understand how the sequence properties of an RNA binding domain and flanking disordered region enable them to cooperate to bind nucleic acids and achieve specific binding affinities. Our findings demonstrate that the ability of the SARS-CoV-2 nucleocapsid protein NTD to bind ssRNA is determined by a combination of sequence composition and the specific positioning of positively charged amino acids within its linear sequence. We identified critical ‘hotspots’ of protein-nucleic acid interaction in the SARS-CoV-2 NTD, where maintaining positive charge allows retention of wild-type binding affinity. These ‘hotspots’ result in a distinctive mode of ssRNA binding in the SARS-CoV-2 NTD-RBD complex, which we observed to be conserved across several coronavirus orthologs, despite significant variations in the NTD sequence. Our study highlights that disordered regions can exhibit conserved interaction mechanisms, even in the absence of exact sequence conservation.

## Results

### “Inert” Intrinsically Disordered Regions Suppress RNA Binding

Our previous work used coarse-grained MD simulations paired with smFRET-based RNA binding experiments to characterize the ability of the SCO2 NTD-RBD to bind ssRNA (52). Simulations and experiments showed that the addition of the disordered NTD_SCO2_ to the folded RBD resulted in a 30-fold increase in the binding affinity for (rU)_25_ compared to the RBD alone. We hypothesized that specific residues in the NTD_SCO2_ formed favorable interactions with RNA, driving the enhanced binding affinity observed. We further speculated that substituting the NTD_SCO2_ with an inert IDR that interacts negligibly with RNA would result in a binding affinity similar to that of the RBD alone. To our surprise, our simulations showed this was not the case.

In the Mpipi model, glycine and serine residues have negligible interactions with RNA or other amino acids. This is in good agreement with prior experimental work that suggests GS-repeat sequences behave as relatively inert Gaussian chains (55, 56). Therefore, we replaced the NTD_SCO2_ with a length-matched GS repeat – (GS)_25_ – and performed simulations with this (GS)_25_-RBD_SCO2_ chimera (**Fig. 2A**). (23, 24, 57). Our simulations revealed repeated association and dissociation events between (rU)_25_ and the (GS)_25_-RBD constructs (**Fig. 2B**), enabling us to calculate an apparent binding association constant, K_A_ (see Methods for details).

**Figure 2.**
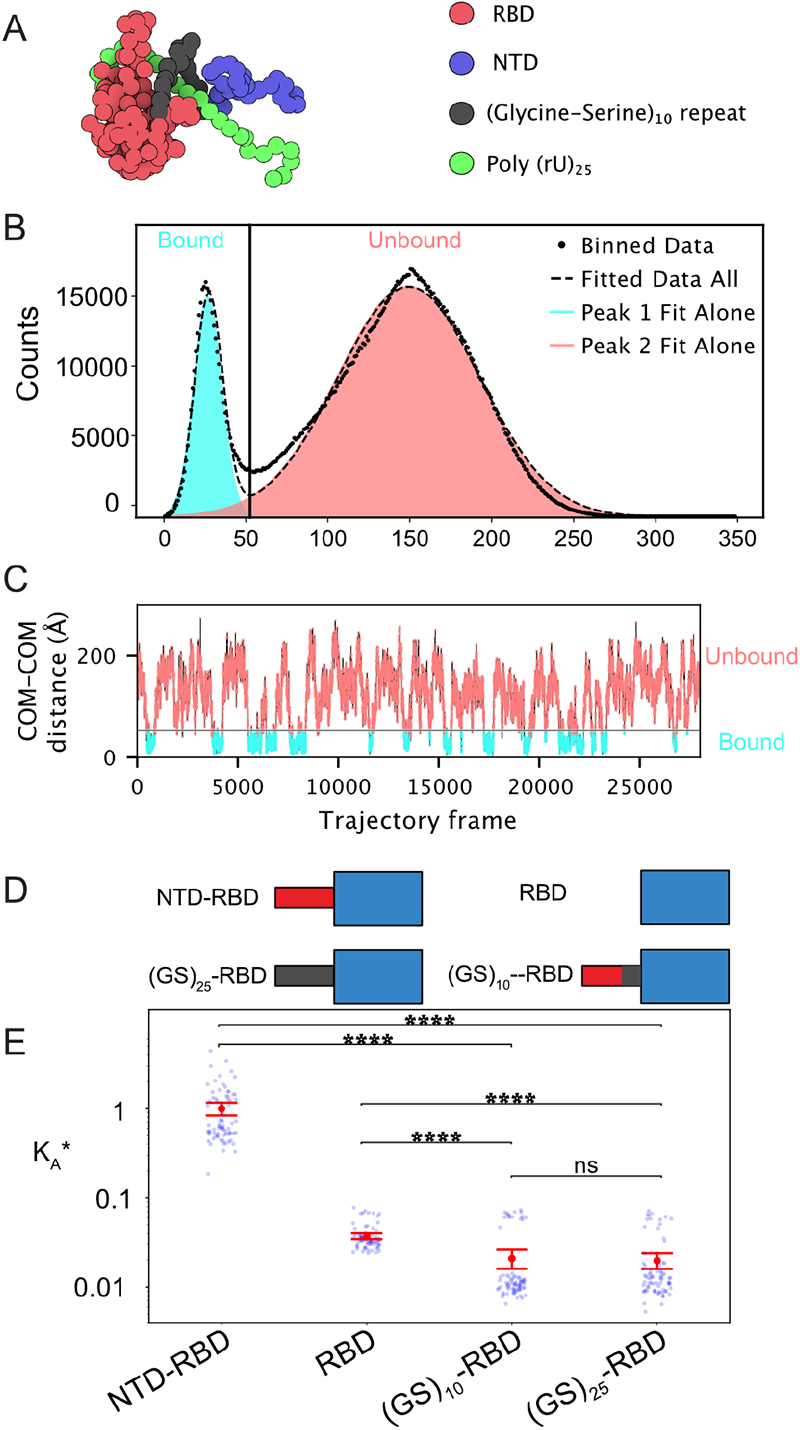
An inert disordered region can suppress a folded domain’s RNA binding ability. **A.** A snapshot of the bound state from a (GS)_25_-RBD + (rU)_25_ simulation trajectory. Simulations utilize the Mpipi forcefield (63). The model represents both amino acids and nucleotides as single beads with specific amino acid-amino acid and amino acid-nucleotide interactions. Folded domains are rigid, and both disordered regions and nucleic acids are dynamic. **B.** The distances between the COM of the (GS)_25_-RBD and (rU)_25_ are plotted over the course of the simulation. A distance threshold (black line) is determined in C (see also Methods) and plotted to delineate the bound and unbound frames. **C.** COM-COM distances from B are plotted as a histogram and show a bimodal distribution that correlates with the bound and unbound states of the protein. The distributions are fitted with dual Gaussians. A distance threshold, which separates bound and unbound frames, is determined by minimizing the overlap of the two populations**. D.** Schematic of the four constructs shown in current “D” + (rU)_25_. **E.** An apparent binding affinity (K_A_) is calculated by utilizing the fraction of bound and unbound frames and Eq. 1. This is then converted to a relative apparent binding affinity (K_A_*) by normalizing all values by dividing by the K_A_ calculated from the SCO2 NTD-RBD + (rU)_25_ simulations. Blue points represent each individual simulation K_A_*, while the red point is the mean of all of the replicate simulations for a given construct. The error bars are the ratio propagated standard error of the mean calculated using Eq. 2. Significance is determined by a Mann-Whitney-Wilcoxon test two-sided with Bonferroni correction. p-value annotation legend: (ns: 5.00e-02 < p <= 1.00e+00), (*: 1.00e-02 < p <= 5.00e-02), (**: 1.00e-03 < p <= 1.00e-02), (***: 1.00e-04 < p <= 1.00e-03), (****: p <= 1.00e-04)

To our surprise, the (GS)_25_-RBD suppressed RNA binding compared to the RBD alone ((GS)_25_-RBD K_A_* = (2.0 ± 0.3) x 10^-2^, whereas RBD K_A_* = (3.7 ± 0.4) x 10^-2^)) (**Fig. 2D**). This result is driven by an entropic effect, whereby the (GS)_25_ impedes the ability of RNA molecules to interact with the RBD. The NTD, in contrast, possesses sequence features that enable direct interaction with RNA, notably residues 30-50, which enhance the macroscopic binding affinity (52, 58).

To investigate the contribution of a smaller and targeted inert region, we next replaced the NTD_SCO2_ 30-50 residue region with a (GS)_10_ linker. While we anticipated a decrease in binding affinity, we expected it to still be stronger than that of the RBD alone. In actuality, we again observed a suppression of RNA binding affinity, with the (GS)_10_ demonstrating weaker binding affinity than the RBD alone, with a K_A_* = (2.1 ± 0.3) x 10^-2^ (**Fig. 2D**), but similar to the (GS)_25_ replaced NTD_SCO2_.

It is widely known that sequence composition and patterning govern the properties adopted by intrinsically disordered regions. However, for IDRs adjacent to RNA binding domains and their binding interfaces, our results suggest that sequence properties can either enhance or suppress RNA binding affinity, depending on the specific IDR sequence. Taken together, our results suggest that the sequence of the N-terminal IDR adjacent to coronavirus RBDs needs to be relatively specific and is most likely conserved, albeit not in the traditional sense of direct sequence alignment; otherwise, without specific residues, the IDR could interfere with RNA binding to the extent of suppressing binding affinity.

### Coronavirus Nucleocapsid Protein NTDs have Conserved Sequence Composition

While NTD’s in coronavirus nucleocapsid proteins appear to always be disordered, their absolute sequence conservation is poor (**Fig. 1B, Supplementary Fig. 3**). If NTDs exist to enhance RNA binding affinity, and disordered NTDs can suppress RNA binding if the ‘wrong’ sequence is present, then how do coronavirus NTDs ensure tight RNA binding is conserved despite largescale variation in sequence?

The decrease in binding affinity caused by (GS)_10_ and (GS)_25_ mutant NTDs indicates that any enhancement in RNA binding provided by the NTD_SCO2_ is sequence dependent. This conclusion is consistent with our prior work, in which small changes in NTD sequence had measurable effects on RNA binding affinity as measured both by single-molecule experiments and by simulations (52).

Operating under the assumption that the NTD_SCO2_ has a role in enhancing RNA binding affinity of the RBD (**Supplementary Fig. 4)**, we reasoned there may be some selective pressure towards NTD sequences that result in a consistent macroscopic RNA binding affinity for the NTD-RBD. Additionally, while RBD structures are highly conserved across coronaviruses, their charged surface residues vary (**Fig. 1D**) (59). As such, we also wondered if there may be a co-evolutionary coupling between the NTD sequence and the RBD surface. Thus despite diverging surface charge of the RBDs, conserved interactions between the NTDs and their respective RBDs could lead to a consistent macroscopic RNA binding affinity.

To investigate this hypothesis, in addition to the NTD-RBD taken from SARS-CoV-2 (SCO2)), we examined NTD-RBD constructs from five other coronaviruses: human coronaviruses OC43, HKU1, and 229E, the Middle East Respiratory Syndrome Coronavirus (MERS), and the Mouse Hepatitis Virus (MHV1). We reasoned that focusing on coronaviruses that predominantly infect the same host would ensure host selective pressures are consistent, thereby minimizing this as a confounding factor to explain differences in RNA binding affinities.

We first examined NTD physicochemical properties that are routinely used to describe IDRs (**Supplementary Table S3-S6**). Despite the large variation in NTD length, all NTDs possess a net positive charge, with the least positive NTD possessing a net charge per residue of +0.056. Expanding this analysis to 45 different coronavirus NTDs, we found no examples in which the net charge was lower than +0.056 (**Supplementary Fig. 5**). This is consistent with RNA binding proteins typically binding RNA through positive electrostatic surfaces that interact with negatively charged RNA (60).

Next, we examined solvent-accessible residues on the RBD surface. We generated five RBD structures for each of the coronaviruses using AlphaFold2, and then took the average of our calculated properties across the five structures (57). The net charge per residue (NCPR) of the RBD surface residues stratified into three categories: relatively positively charged (229E = 0.126, SCO2 =0.066, MERS = 0.052,) neutral (HKU1 = 0.0, MHV1 = -0.011), and negatively charged (OC43 = -0.053).

In summary, while the surface charge of the RBD domains appears more variable, our analysis suggests an extremely strong bias for coronavirus NTDs to retain a net positive charge, in line with our expectation that these IDRs facilitate enhanced RNA binding. Compositional conservation in the NTD, or the retention of specific physicochemical features (such as net charge), could enable conserved interactions, despite lack of absolute sequence conservation.

### Sequence Composition Alone Does Not Determine NTD Contribution to Binding Affinity

Since NTD composition is relatively consistent across orthologs, we wondered if sequence composition alone was sufficient to dictate RNA binding affinity. To test this, we performed a tiling experiment. Here, we repositioned the previously identified 30-50 residue positive charge block region of the SCO2 NTD (NTD_SCO2_) that we and others showed to be involved in single-stranded RNA binding (52, 58). We placed this charge block at positions 1, 6, 11, 16, 21, 26 (referred to as mutants T1, T6, T11, T16, T21, T26) and 31 (wild type) of the NTD_SCO2_ (**Fig. 3A**). Finally, we calculated apparent binding affinities of each of these variants with (rU)_25_. These sequences maintain the same sequence composition but rearrange the amino acids, which allows us to determine whether there are positional contributions to RNA binding or if sequence composition alone is sufficient to achieve RNA binding.

**Figure 3.**
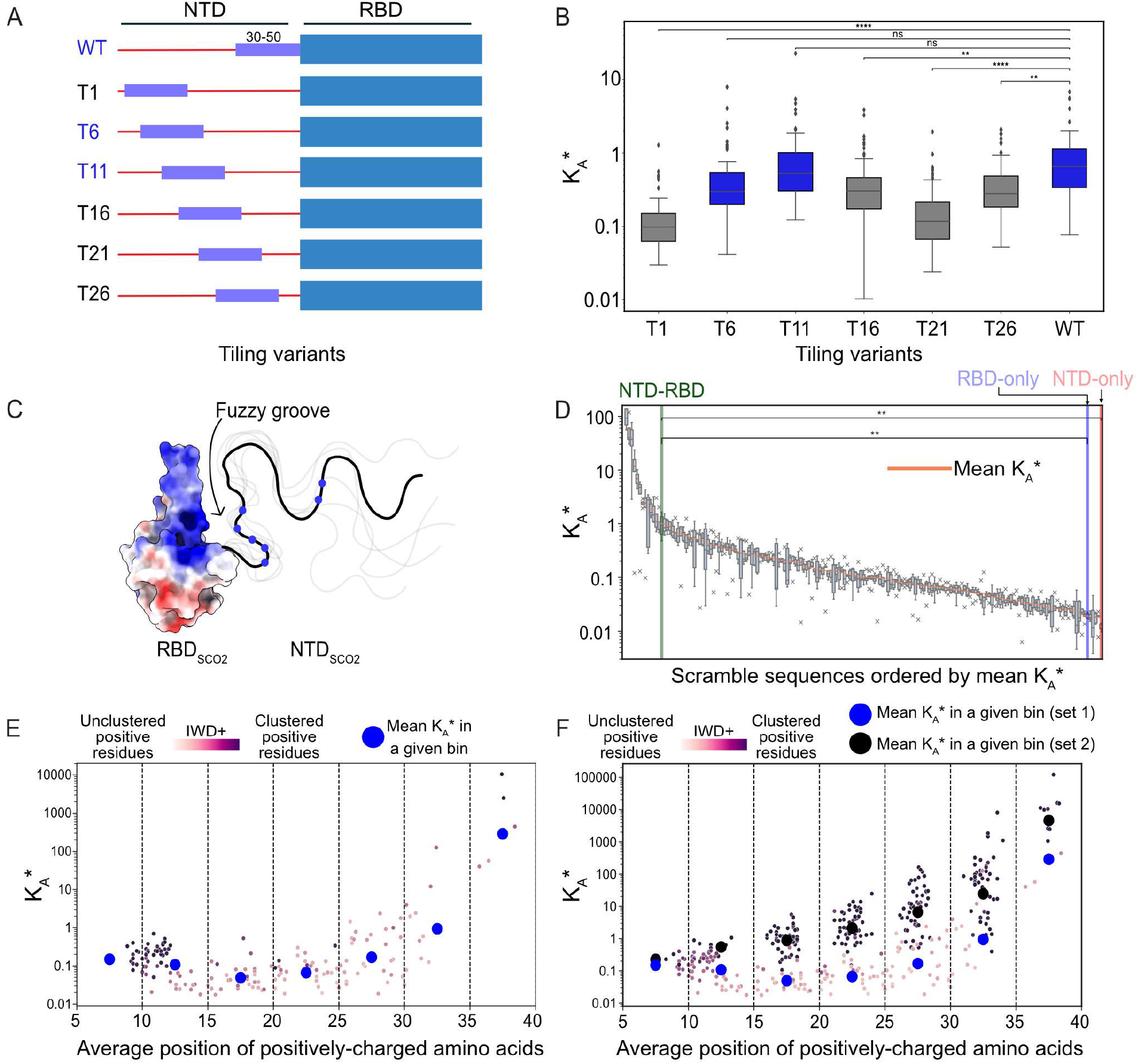
Clusters of positively charged residues determine the affinity enhancement provided by the NTD on RNA binding. **A.** Schematic showing the wild type and tiling mutants that systematically reposition residues 30-50 from the wild-type sequence. **B.** Binding affinity for tiling mutants schematized in panel A. Tiling mutant T6 and T11 show wildtype-like binding affinity, whereas all other variants show binding affinity less than the wild type. **C.** Graphical schematic highlighting the positively-charged “fuzzy groove” that can form upon RNA binding between the positively-charged beta extension on the RBD and the cluster of positively charged residues on the NTD. In the RBD positively charged surfaces are colored blue, negatively charged surfaces are colored red, and neutral surfaces are colored white. A representative NTD is drawn with the blue circles representing the relative positions of the positively charged residues. **D.** Binding affinities for 172 scramble variants. Each variant reports on the binding affinity for an NTD-RBD construct, where for each variant the NTD sequence was randomly scrambled. Despite having an identical amino acid composition, sequence order enables a four-order-of-magnitude change in binding affinity, highlighting the importance of sequence in dictating binding affinity. **E.** Scramble sequences plotted with binding affinity vs. the average position of positively charged residues distributed across the sequence. For positional bins, average binding affinity is shown as a blue circle. Individual points are colored based on the IWD+ score, which reports on the clustering of positively charged residues (darker colors = more highly clustered). **F.** Same data as shown in E, with an additional set of scrambles designed to cluster positively charged residues. The average binding affinity of this second set is shown as black circles.

To our surprise, the relative position of positive tiles has a significant impact on the apparent binding affinity (**Fig. 3B**). Two mutants showed wild type-like binding affinities, yet the others bound RNA more weakly. This suggests that the relative location of positive charge with respect to the RBD tunes RNA binding affinity. Simultaneously, this result lends credence to a model in which mere sequence composition is not sufficient to achieve ‘adequate’ (wild type) binding affinity.

To further test how sequence composition impacts RNA binding, we generated 386 scrambled NTD_SCO2_ sequences in which the sequence composition is identical, yet the order of the amino acids has been changed. An initial set of 172 scrambles were generated in four ways: The first by randomly shuffling the NTD_SCO2_; the second by shuffling the NTD_SCO2_ while also making each amino acid change be as chemically different from the wild-type sequence as possible in terms of charge and aromaticity; third, by shuffling the NTD_SCO2_ while forcing positively charged residues from falling in the 30-50 residue region; and fourth, by shuffling the NTD_SCO2_ while restricting the majority of charged residues to the 30-50 region or a region spanning residues 4-17. Using these scrambled sequences, we performed coarse-grained MD simulations to measure K * with (rU) .

Binding affinities were calculated for each of the scrambled sequences and compared with one another (**Fig 3D, Supp. Table 7**). The dynamic range of K * observed here spans five orders of magnitude, demonstrating the dramatic impact relative amino acid position can have on binding affinity. However, for the majority of the scrambled sequences, the binding affinity is fairly similar, and, importantly, this “average” binding affinity is almost an order of magnitude weaker than the wild-type NTD-RBD.

Taken together with our tiling simulations, these results suggest composition is not the sole determinant of how the NTD_SCO2_ influences RNA binding. While 172 scrambled sequences is only a fraction of the total number of possible sequence compositions that could be generated for the NTD_SCO2,_ the observation that the wild-type NTD_SCO2_ sequence is among those with the highest apparent affinity suggests that the ordering of the residues in the NTD_SCO2_ is specific.

### Disordered Region Residue Sequence Positioning Dictates RNA Binding Capacity

While most scrambled sequences had similar binding affinities that were much weaker than the wild-type sequence, we identified a subset of sequences that had binding affinities equal to or greater than that of the wild-type sequence. Based on our tiling simulations, we reasoned that the relative position of positively charged residues might underlie the increased binding affinity of these select sequences, highlighting regions of the NTD that are more binding-competent.

To assess how the position of positively charged residues correlates with binding affinity, we plotted binding affinity versus the average position of all positively charged residues in each scrambled sequence that we initially tested (**Fig 3E**, blue circles are the binned means of the first 172 sequence). The average position is calculated as the mean of the location of the arginine and lysine residues in the linear sequence of the NTD_SCO2_. This analysis revealed a correlation between strong binders and the average position of positively charged residues. When the average position of positive residues is around residues 30-40, binding affinity is drastically increased in comparison to the other regions. This same region is relatively positively charged in the wild-type NTD_SCO2_.

The importance of the position of positively charged residues offers a ‘structural’ explanation for the enhanced binding affinity afforded by the wild-type NTD. Charged residues within this region enable the formation of a ‘fuzzy groove’. One half of this groove is made of the positively-charged RBD, while the other half comes from the NTD. This fuzzy groove enables simultaneous interactions between the NTD_SCO2_ and the RBD_SCO2_ with RNA and, thus, tight RNA binding (**Fig 3C**).

Curiously, we observed a relationship between the average positioning of positively charged residues and binding affinity with similarities to our tiling simulations. We binned the scrambled sequences by average positive charge positioning and compared their binding affinities. The two bins that spanned residues 5-10 and 10-15, had binding affinities on average equal to the bin that contained the wild-type sequence. Bins that spanned residues 15-20 and 20-25 were each significantly different from the wild type bin (p = 0.00013 and 0.016, respectively), and both were weaker on average than the wild type bin. Regions that clustered their charge between residues 30-35 and 35-40 had significantly higher binding affinities in comparison to the wild type bin. This supports our hypothesis that the relative positioning of positive residues greatly influences the binding affinity and that certain NTD regions are more binding-competent than others.

We next investigated how the arrangement of positively charged residues impacts the variability of binding affinities within each region. Despite observing variations in binding competence among different regions on average, there was still a wide range of affinities within each region. We hypothesized that the clustering of charged residues, which is not captured by averaging their linear positions, influences binding affinity.

To visualize this, we used an inverse weighted distance (IWD+) metric to calculate the charge clustering of the positive residues arginine and lysine. When plotting the IWD+ values over our binned data (**Fig 3E**), we observed relatively consistent positive clustering for most sequences we generated. However, we noticed that the bins spanning residues 5-15 exhibited higher positive clustering due to the N-terminal positioning of the average positive residues. Even within these bins, there were sequences with lower clustering and weaker binding affinities compared to highly clustered sequences. Additionally, the wild-type sequence showed higher positive clustering and had an increased binding affinity compared to sequences with less clustering within the region encompassing residues 25-30.

We noticed several sequences in different regions that exhibited significantly increased clustering of charged residues and binding affinities. We then created a second set of sequences. Our aim was to enhance the average binding affinity in each region by clustering the charged residues. Given the generally higher binding affinities for sequences with positively charged residues clustered closer to the 30-50 amino acid region, we expected to observe a response where highly clustered sequences would have increased binding affinities in comparison to lower clustered sequences. We also expected that the average positioning of positively charged amino acids, when closer to the C-terminal end of the NTD, would have higher binding affinities than sequences with charged residues positioned closer to the N-terminal portion of the NTD.

To test this hypothesis, we generated sequences by scrambling and then constraining the final sequences to have all seven positively charged residues within ±2 residues of their respective bin boundaries. This resulted in sequences with increased positive clustering, as indicated by the IWD+ metric. We calculated the binding affinity of these sequences, using the same methodology as the initial scrambled sequences, plotted them alongside their IWD+ values, and compared them to the original sequences (**Fig 3F**). As anticipated, the highly clustered sequences exhibited, on average, increased binding affinities in each region. We classified the sequences based on either their clustering similarity to the wild-type sequence or significantly higher clustering. Sequences with clustering similar to the wild type followed the previous tiling experiments in terms of how sequence location affected binding affinity. On the other hand, sequences with increased clustered charge showed higher binding affinities, often comparable to the wild-type sequence. Further, the clustered positively charged sequences displayed an exponential relationship between the proximity to the C-terminal region and their apparent binding affinity, highlighting how the positioning of charge impacts NTD-RBD ssRNA binding.

Analysis of the bound-state trajectories revealed a dynamic or “fuzzy” complex in which specific subregions of the NTD_SCO2_ contact the RNA. From the simulations of scrambled NTD_SCO2_, we observed that particular regions of charge are sufficient to increase binding affinity. Thus, we hypothesized that we would find such charged patches within orthologous NTD sequences that exhibited increased binding affinity. Here we propose a model for conserved disorder without conserved linear sequence. Similar to how IDRs that contain SLiMs can exhibit sequence heterogeneity as long as short motifs are maintained(61), regions that contain conserved charge clustering can also have high sequence dissimilarity but still maintain sufficiently strong binding affinity for ssRNA.

### NTD-RBD:RNA Behavior in the Bound State is Conserved Across Orthologs

Our scrambles confirm that the NTD sequence has a substantial impact on NTD-RBD RNA binding affinity. We therefore asked if natural NTD sequences encode a similar “fuzzy groove” binding mode, despite seemingly large-scale variation in NTD sequence and RBD surface chemistry. In this model, specific subregions of the NTD come into closer proximity to the RBD driven by favorable NTD-RNA interactions on one side and RBD-RNA interactions on the other (**Fig. 3C**). To test this, we performed simulations of each of the six ortholog NTD-RBD constructs with (rU)_25_ and assessed the bound-state conformational ensemble of the NTD.

Bound-state ensembles were visualized using scaling maps. Scaling maps capture the average inter-residue distance between all pairs of residues for RNA-bound conformers. We normalized the scaling maps by the inter-residue distance of sequence-matched NTD-RBD simulations performed in the absence of RNA (**Fig. 4A**). Shades of blue reflect distances that are closer together in the bound state, while shades of red denote regions that are further apart in the bound state. For SCO2, this analysis identified two regions in the NTD that are closer to the RBD in the bound state ensemble centered around residues 10-20 and residues 30-50, similar to our tiling simulations and as reported previously(52). This analysis can be done selectively for one of the residues in the NTD to visualize where it increases RBD interactions when bound to RNA by mapping its distances across the entire NTD-RBD construct with RBD residues colored with respect to NTD distance (**Fig. 4B**). Doing so shows that in the bound state, the NTD moves closer to the positively charged RBD β3 extension, highlighting the formation of a fuzzy positive groove between the positive β3 extension and positive subregions in the NTD_SCO2_.

**Figure 4.**
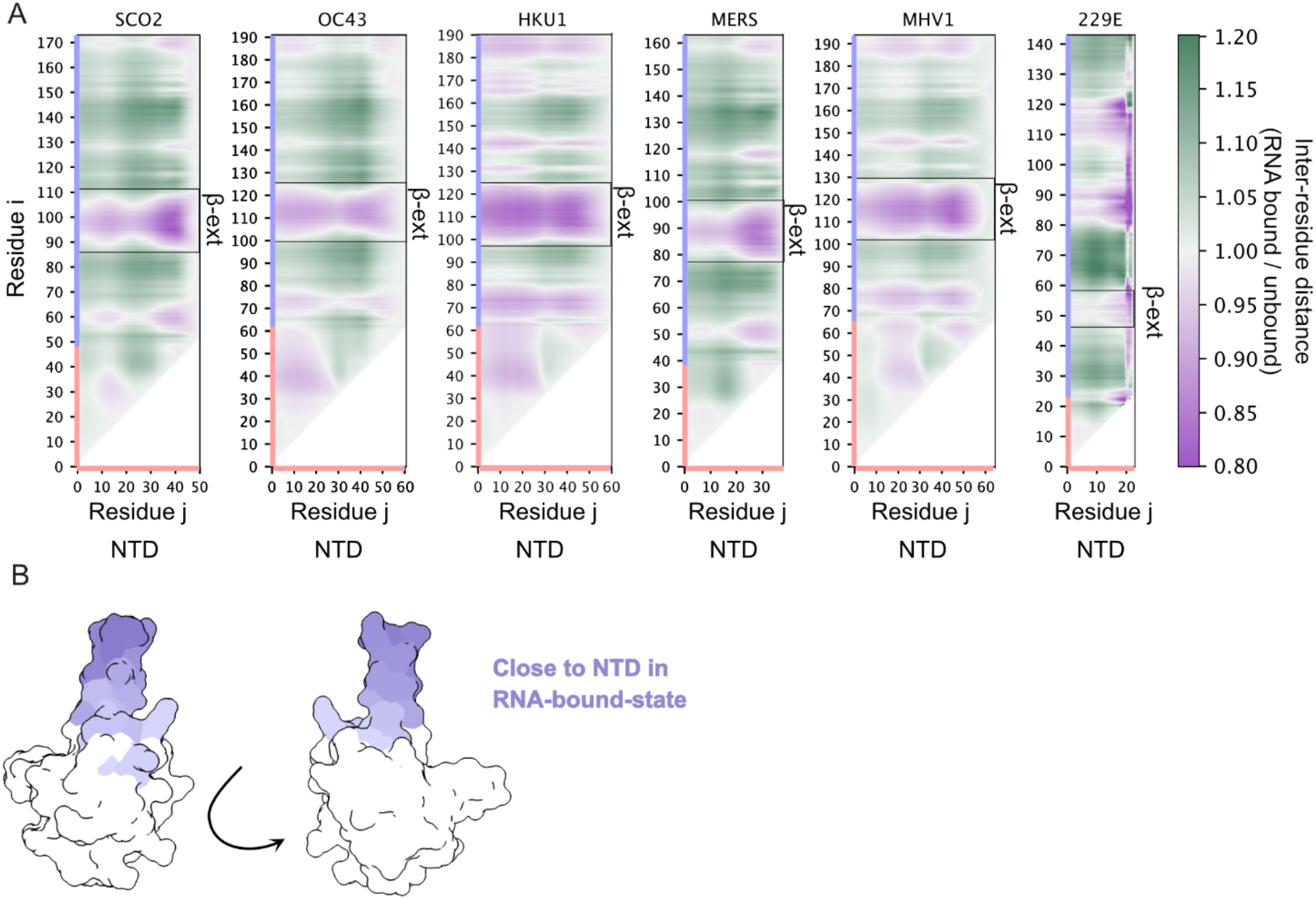
Orthologous nucleocapsid proteins show similar bound-state ensembles despite variations in RBD surface charge residues and NTD sequence. **A.** Scaling maps quantify the average inter-residue distance between NTD residues (X-axis, colored pink) and NTD or RBD residues (Y-axis, colored pink and light blue respectively) in the bound state. Heatmap values are calculated by calculating the average inter-residue distance in the RNA-bound state and dividing that distance by the average inter-residue distance in the RNA-unbound state. Purple colors report on inter-residue distances that are closer together in the bound state while green colors report on inter-residue distances that are further apart in the unbound state. In all six orthologs, the NTD is closer to the β-extension in the bound state, reporting on the formation of a positive fuzzy groove in the bound state. **B.** Regions closer to the NTD in the RNA-bound state are highlighted on the SCO2 RBD structure in shades of purple with more intense purple signifying closer on average.

We repeated this analysis for the remaining five orthologs to determine if these NTDs also move closer to the RBD. In line with our expectations, this analysis reveals that in all cases, two specific subregions within the NTD come closer to the RBD. Despite large-scale variation in both folded-domain surface charge and NTD sequence, the mode of RNA binding appears to be largely conserved across the six coronavirus NTD-RBD constructs examined.

## Discussion and Conclusion

Intrinsically disordered proteins and protein regions are prevalent across eukaryotic, prokaryotic, and viral proteomes. They play a wide variety of essential roles yet – perhaps paradoxically – often appear to be relatively poorly conserved sequences by alignment. In this study, we sought to understand how a specific molecular function (RNA binding) could be conserved despite large-scale changes in amino acid sequence. We utilized two domains of various coronavirus nucleocapsid protein orthologs as a convenient model that contains both a disordered region (NTD) and a folded domain (RBD) that binds RNA. Despite poor sequence conservation assessed by alignment across NTDs, we found that the orthologs were compositionally conserved. That is, the orthologs have similar charge properties in both the NTD and portions of the RBD. Specifically, NTDs harbor a net positive charge, while RBDs retain specific positively charged regions on their surface. Despite this conservation, the length and sequence of N protein NTDs vary dramatically, and while RBDs maintain the same 3D structure, orthologous RBDs showed a diverse set of surface properties, from highly negatively charged to highly positively charged.

To assess how the sequence composition of the disordered NTDs influences interactions with the RBDs and impacts RNA binding, we performed various coarse-grained molecular dynamics simulations of coronavirus nucleocapsid proteins with single-stranded RNA. These simulations enabled us to interrogate the role of sequence composition and residue positioning in coronavirus NTDs ability to increase binding affinity of the NTD-RBD. We first showed, that replacing the

NTD with a glycine-serine repeat sequence suppresses RNA binding, illustrating the impact that IDR sequence can have on intermolecular interactions. By testing hundreds of different sequences with the same overall composition, we determined that composition alone does not dictate RNA binding affinity. Instead, our simulations highlight the importance of clusters of positively charged residues, and that the relative position of positive clusters along the NTD also matter. Taken together, our use of rationally designed synthetic sequences illustrates that, at least in simulations, the absolute linear sequence can have a profound impact on IDR-mediated molecular interactions, even for simple systems using simple physics-based models.

Finally, we performed simulations of five orthologous NTD-RBD constructs, noting that despite dramatic changes in both the charge properties of the RBDs and the NTD sequence, the bound-state ensemble conformational properties were conserved, with the NTD wrapping against the positively charged beta-extension, forming a so-called “fuzzy groove” that can accommodate RNA. Despite differing sequences, we uncovered a similar mode of interaction between the NTDs and their RBDs, with each NTD having two ‘hotspots’ of interaction that coordinate to interact more often in the bound state with their RBD’s positively charged β3 extension. Curiously, the ortholog for which these hotspots are least prominent (229E) also has the most positively charged RBD, pointing to a potential mechanism to compensate for a ‘weaker’ (less positively charged) NTD. Both our tiling simulations and scramble simulations of the NTD_SCO2_, showed positional contributions to RNA binding affinity, which corroborated the role of the ‘hotspots’ in increasing binding affinity and supported a model where proper positioning of specific residues, in this case positively charged, is the important factor for a flanking disordered NTDs ability to increase RNA binding affinity. While they lack absolute sequence conservation, the conserved nature of these hotspots and their interactions with the RBD β3 extension opens up the possibility of developing inhibitors that can interact preferentially with the β3 extension to modulate the nucleocapsid proteins ability to bind ssRNA. The conservation of ensemble conformational properties in the absence of sequence conservation highlights how IDRs can simultaneously facilitate functional conservation despite supporting highly variable sequences.

While this study focused on the NTD-RBD from coronavirus nucleocapsid proteins, we expect that the information learned here will be widely applicable to a range of disordered nucleic acid-binding proteins. While absolute sequence conservation may not be present, there is still the possibility of conserved behavior encoded into diverging sequences. Rather than solely focusing on sequence alignments to provide information on conservation and important residues, quantitatively describing the ensemble that a disordered region takes on and assessing how it behaves with and without its ligand(s) may provide better insight into the residues that are important and sequence features that need to be maintained to ensure proper biological function.

## Methods

### Molecular Dynamics Simulations

All simulations were performed using the LAMMPS simulation engine(62). We performed molecular dynamics simulations in the NVT ensemble using the default parameters of the physics-driven coarse-grained force-field Mpipi developed by Joseph et al. (63) The model represents both amino acid residues and nucleotides as chemically unique singular beads and was parameterized to recapitulate the behavior of disordered proteins in isolation as well as their ability to undergo phase separation with and without RNA. Inter-bead interactions consist of a combination of short-range contributions from a Wang-Frenkel potential, which captures a combination of Van der Waals, cation-pi, and pi-pi interactions, and a long-range Coulombic potential for amino acids with net charge and RNA nucleotides. The ability of the Mpipi force field to recapitulate disordered protein dimensions has been previously shown (63, 64). Simulations were performed under an effective ionic strength of 50 mM NaCl, conditions we previously found to engender good agreement between simulation and experiment when comparing with experimentally-measured RNA binding affinities using single-molecule experiments (52).

We also assessed the ability of the Mpipi forcefield to recapitulate single-stranded RNA (ssRNA) dimensions by comparing simulations of (rU)_40_ with scattering data from small-angle X-ray (SAXS) experiments for the same construct(65). This comparison revealed excellent agreement across the full scattering curve and in terms of the scattering-derived radius of gyration; using the Molecular Form Factor approach of Riback et al., R_g_ ^sim^ = 30.9 ± 0.1 Å while R_g_ ^exp^ = 30.2 ± 0.3 Å) (**Supplementary Fig. 1**) (66).

Simulations were performed in a 30 nm^3^ simulation box with periodic boundary conditions. Protein and RNA are allowed to diffuse freely throughout the box. Disordered regions and ssRNA behave as dynamic flexible polymers, sampling an ensemble of conformations (63). However, as done previously, folded domains were made rigid, and residues buried within folded domains experienced downscaled non-bonded interactions(52, 63). Unless otherwise specified, all simulations were run for 300 million steps per replicate. The exceptions are the ‘scrambled’ simulations, which were run for 100 million steps per replicate. Protein and RNA configurations were saved every 10,000 steps, and the first 0.2% was removed for equilibration. Visualization of protein-RNA complexes was done with Protein Imager and VMD (67, 68). Simulations were analyzed using SOURSOP and MDTraj (69, 70). Small angle X-ray scattering was analyzed using the Molecular Form Factor (MFF) (http://sosnick.uchicago.edu/SAXSonIDPs), while synthetic scattering data for simulations were generated using FOXS default settings(66, 71).

We performed simulations of the NTD-RBD, NTD, and RBD of six coronavirus orthologs. Specifically, we examined five coronaviruses that infect humans: SARS-CoV-2 (SCO2), Middle Eastern Respiratory Syndrome virus (MERS), Human Coronaviruses OC43, Human Coronavirus HKU1, and Human Coronavirus 229E, as well as Murine Hepatitis Virus (MHV1). Sequence alignments were compared to determine a region of the RBD that was well conserved between all orthologs to delineate the start and end positions of the NTD and RBD’s of each ortholog (54, 72–74). For simulations with ssRNA, all simulations were done using (rU)_25._

To capture conformational heterogeneity in an artificially rigid structure, we utilized Colabfold to generate five different starting structures for each coronavirus orthologous RBD (23, 57). For simulations of wild-type versions of each ortholog’s NTD-RBD all five starting structures are used, to enable conclusions to be less biased by a specific starting conformation. As expected, certain RBD conformers bind RNA better than others, but in all cases where different NTDs are compared, the same sets of RBD conformers are used, such that any RBD conformation-specific biases are consistent across the set (**Supplementary Fig. 2**). For the large scrambled library, 1 conformation for the SCO2 RBD is used. All simulations were run with multiple replicates per starting RBD structure, with a minimum of five replicates per RBD conformation.

### Limitations of Coarse-Grained Simulations

Our use of the Mpipi model should not be taken to imply that RNA or proteins are faithfully represented at one bead per residue/nucleotide resolution. Both proteins and RNA are complex biomolecules with many degrees of freedom, a chemically heterogeneous structure, and can engage in a variety of sequence and structure-specific interactions that are not captured by a simplified coarse-grain model. Our goal in using a simplified coarse-grain model is to enable high-throughput biophysical assessment in a system that, based on prior work, we have good reason to believe is semi-quantitative in terms of relative accuracy (52, 63). While we refer to the molecules in our simulations as protein and RNA, in reality, they are better thought of as RNA- and protein-flavored polymers. The simplicity of this model enables us to address questions that would be intractable using either higher-resolution simulation approaches or experiments. Despite this, we are under no illusion regarding the simplifying assumptions made for a coarse-grain model.

### Calculating Apparent Association Constants From Simulations

We determined apparent association constants (K_A_) by using an updated version of our previous center of mass (COM) calculations that were able to qualitatively recapitulate SARS-CoV-2 NTD-RBD single-stranded RNA binding (52). To do this, post-equilibration simulation frames were divided into bound and unbound states. This delineation was achieved by first taking the intermolecular center-of-mass distances between the protein and the RNA and plotting the distribution of distances. The histogram of intermolecular distances follows a bimodal distribution that reports on the bound and unbound states, and can be fit with two Gaussians (**Fig. 2C**). We then determined the intersection that minimizes the overlap of the two distributions to define a cutoff distance. The cutoff distance varies based on the size of the protein and RNA. Finally, as done previously, we classify frames as bound or unbound by assessing the linear intermolecular COM distance trajectory and delineating frames as bound when five or more frames are below the cutoff distance. This minimum number of consecutive frames allows us to distinguish between transient random interactions between protein and RNA vs. encounters with a reasonable “lifetime”, implying direct and continuous interaction. The distributions and distance cutoffs are calculated for every set of NTD_a_-RBD_b_ + (rU)_n_ simulations, where *a* and *b* represent specific NTD or RBD sequences and *n* the length of the single-stranded (rU), allowing us to determine protein-RNA specific distance thresholds for each simulation.

The resultant fraction of bound frames is used to calculate an apparent K_D_ with the equation:

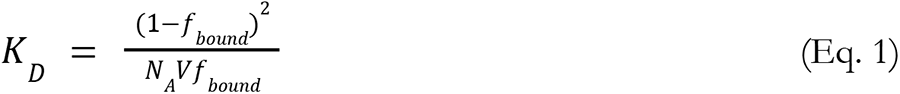

Here *f_bound_* refers to the fraction of frames where the protein and RNA are determined to be in the bound state from our COM-COM distribution analysis. *N_A_* refers to Avagodro’s constant, and V is the simulation box volume in liters, which returns a *K_D_* in mol/L. *K_A_* is then calculated using the xpression *K_A_* = 1/*K_D_* . While we determine if two molecules are bound or unbound in a different manner, this approach is analogous to that of Tesei *et al*. (75).

It is important to note that the K_A_s determined from these simulations are not meant to represent absolute values that would be comparable to those determined from experiment. Our prior work has shown that K_A_s calculated from Mpipi simulations for this system lack absolute agreement with experimentally measured values. Despite this, when experiment and simulation-derived K_A_ values are normalized by an internally consistent reference (i.e., the K_A_ obtained from NTD-RBD binding (rU)_25_), we see good agreement between simulations and experiment, both as a function of RNA length and as a function of the presence/absence of the NTD (52). To that end, binding affinity here is reported as K_A_*, a normalized binding affinity we define as the ratio of the apparent K_A_ of a given protein + RNA simulation divided by the corresponding K_A_ for the analogous SCO2 NTD-RBD binding to (rU)_25_. This enables the SCO2 NTD-RBD + (rU)_25_ simulations binding affinity to be a reference point with which to understand the strength of interactions of other orthologs. All K_A_* values are thus greater than 1 (stronger binding than the SCO2 NTD-RBD + (rU)_25_) or less than 1 (weaker binding than the SCO2 NTD-RBD + (rU)_25_).

Error is propagated for our ratio (K_A_*) using:

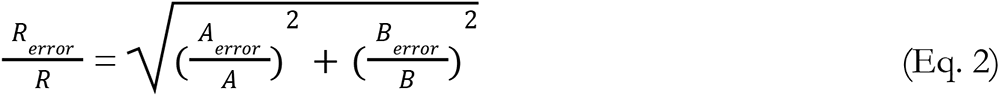

*R and R*_error_ here represent the ratio and the error of the ratio. *A* and *B* represent the numerator and denominator of our ratios, respectively, and A_error_ and B_error_ are their associated errors (standard error of the mean).

### Calculating Charge Clustering in Disordered Regions

Charge clustering is quantified by the inverse weighted distance (IWD), a metric that has been applied to study amino acid clustering in several systems (76–79). Unlike the patterning parameters κ (“kappa”) or sequence charge decoration (SCD), which quantify the patterning of oppositely charged residues with respect to one another, here our interest is on the clustering of positive residues only(80, 81). The IWD score allows us to quantify the clustering of a specific subset of residues. When residues are clustered together, the IWD score is high, whereas when residues are evenly distributed, the IWD score is low. IWD scores were calculated using sparrow (https://github.com/idptools/sparrow).

### Statistical Analysis

Every simulation has a minimum of five independent replicates, and calculated values are presented as 95% confidence intervals (box plots, with medians marked), mean and standard error of the mean, or geometric mean and geometric standard deviation (clarified in text below figures). Fitting of Gaussian distributions was done in Python using scipy.optimize.curve_fit (82).

## Data Availability and Software

Analysis code and data (calculated distance distributions and contact map information) are deposited at https://github.com/holehouse-lab/supportingdata/tree/master/2023/alston_2023. For further information on the use of code, please refer to the deposited Jupyter notebooks.

## Acknowledgments

We thank members of the Soranno lab and Holehouse lab for many useful discussions over the years. We particularly thank Dan Griffith, who has provided useful insights for data visualization with Python. We thank Dr. Emery Usher for help in performing Guinier analysis of (rU)_40_ scattering data. We would like to thank Dr. Jerelle Joseph for the parameterization of the Mpipi force field, which enabled us to do this work. We thank Dr. Lois Pollack and Dr. Steve Meisburger for sharing scattering data for (rU)_40_. Funding for this work was provided by the National Institute of Allergy and Infectious Diseases with R01AI163142 to A.S.H. and A.S., by the Human Frontiers in Science Program (HFSP RGP0015/2022) to A.S.H, and by the National Cancer Institute with an F99CA264413 to J.J.A.

