## Supplementary Information for "Conserved molecular recognition by an intrinsically disordered region in the absence of sequence conservation"

### Supplementary Figures

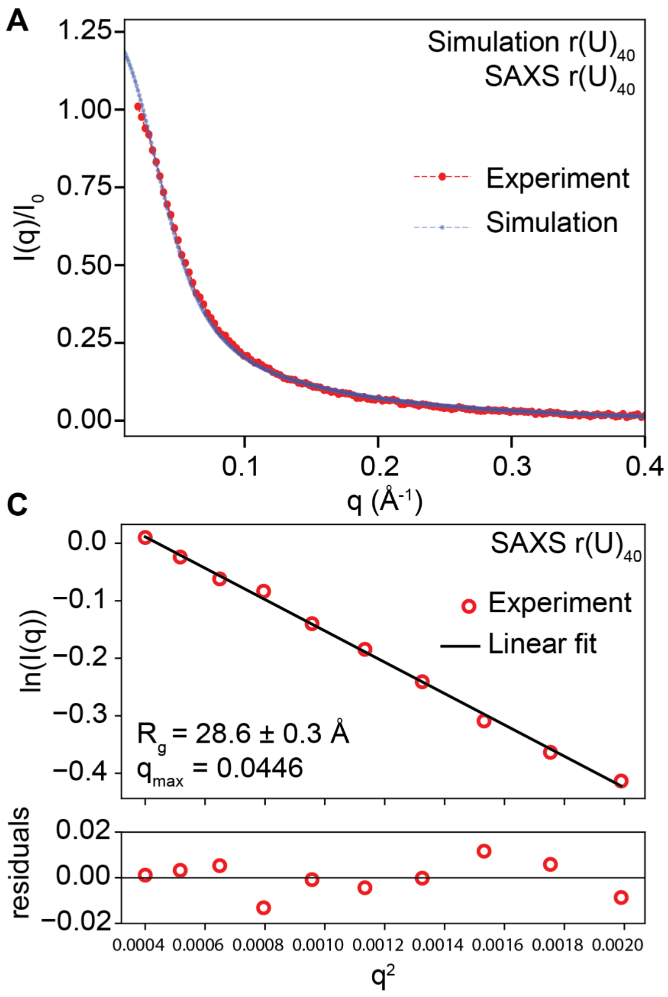


**Supplementary Figure 1. The Mpipi forcefield captures the experimental dimensions of ssRNA. A.** To assess how well rU homopolymeric RNA molecules behave in Mpipi, we compared small-angle X-ray (SAXS) scattering profiles obtained for $(rU)_{40}$ with scattering profiles generated from simulations of $(rU)_{40}$ using FOXS(71). The agreement is extremely good, as shown by the tight overlay of the simulated and experimental scattering curves. **B.** We estimated the $(rU)_{40}$ radius of gyration ($R_g$) using the Molecular Form Factor (MFF) approach of Riback et al. (66). This approach yielded an $R_g$ of 30.2 ± 0.3 Å. Analyzing synthetic scattering data from simulations in the same way yields an $R_g$ of 30.9 ± 0.1 Å, while calculating the Rg directly from simulations gives an Rg of 32.2 Å.

**C.** As a complementary analysis we also analyzed the SAXS data using Guinier analysis, fitting up to $qR_g < 1.3$. Based on this analysis we calculated a slightly smaller $R_g$ of 28.9 ± 0.3, although this value is in good agreement with both simulations and the $R_g$ obtained from fitting to the MFF.

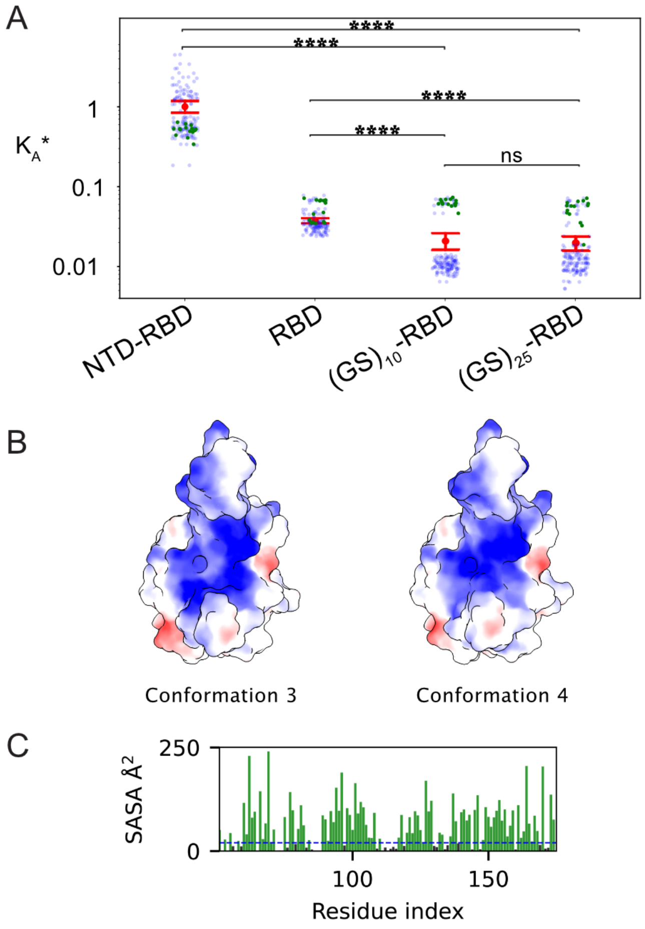


D

| Conformation | NCPR |
|---|---|
| RBD 1 | 0.0659 |
| RBD 2 | 0.0667 |
| RBD 3 | 0.0659 |
| RBD 4 | 0.0682 |
| RBD 5 | 0.0659 |

E

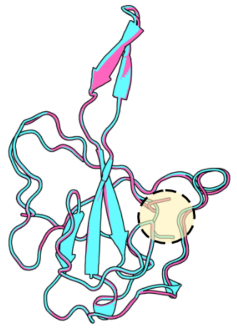

**Supplementary Figure 2. Structural heterogeneity in the RBD impacts the relative apparent binding affinity A.** The relative apparent binding affinity ($K_A$*) plot from Figure 2E is plotted with $RBD_{SCO2}$ conformation four plotted in green, highlighting its average binding affinity differs from the other conformations. For the wildtype NTD-RBD conformation, four behaves similarly to other conformations. However, for the RBD alone and GS mutants, it is a better binder of non-specific

RNA than the other conformations. This can be explained by reasoning that for constructs where the binding affinity is dominated by the RBD alone, the properties of the RBD will greatly affect binding affinity. However, where binding is a combination of NTD and RBD interactions, the affinity will be affected by how the two domains cooperate to bind RNA. To test this hypothesis we examine the charge properties of the different conformations. **B.** Structures of conformation 3 and 4 of $RBD_{SCO2}$ with charge patterning determined by ChimeraX Coulombic electrostatic potential. While small, charge distribution differs around the β-extension that is involved in RNA binding. **C.** Surface-accessible residues are calculated for all $RBD_{SCO2}$ conformations (conformation 4 is shown as an example). Green bars represent residues that are surface accessible, while black bars show residues that are buried. **D.** Net charge per residue (NCPR) is calculated for all surface-accessible residues for each conformation. Conformation 4 has a higher surface-accessible NCPR than the other constructs. **E.** Overlay of conformation 3 (pink) and 4 (teal). Conformation 4 has a shift in its N-terminal residues that alters its accessible charge patterning as highlighted by the beige circle.

```
MHV1: --------------MSFVPGQENAGGRSSSVNRAGNGILKKTTWADQTERGPNNQNRGRRNQPKQTATTQPNSGSVV-
OC43: --------------MSFTPGKQSS-SRASSGNRSGNGIL---KWADQSDQVRNVQTRGRRAQPKQTATSQQPSGGNVV
HKU1: --------------MSYTPGHY-AGSRSSSGNRS--GILKKTSWADQSERNYQTFNRGRKTQPKFTVSTQPQGNTIP-
SCO2: MSDNGPQNQRNAPRITFGGPSDSTG-SNQNGERSGARSK-----------QRR-----PQGLPNNT------------
MERS: --------------------------------MASPAAPRAVSFADNNDITNTNLSRGRGRNPKPRAAP---------
229E: --------------------------------------MATVKWADASEPQRGRQG----------------------
```

**Supplementary Figure 3. Multiple Sequence Alignment of Coronavirus N-Terminal Domains**

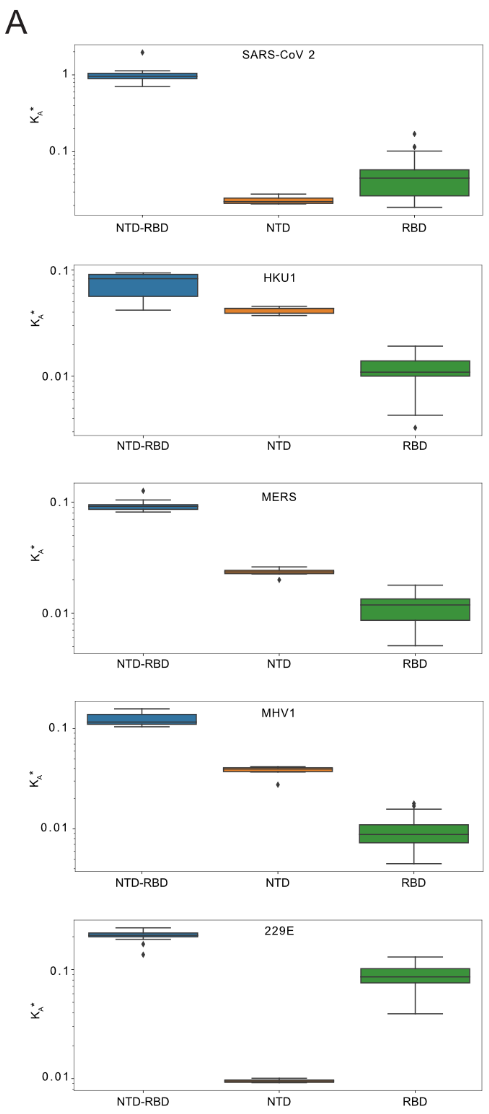


**Supplementary Figure 4. For all NTD-RBD orthologs, the combination of NTD and RBD has an increased binding affinity than RBD alone. A.** Five conformations of each orthologous RBD were generated by Colabfold and simulated with their NTD and $(rU)_{25}$. Relative binding affinities were calculated as stated in the methods for the NTD, RBD, and NTD-RBD simulations. OC43 is not shown due to NTD and RBD binding alone being too weak to fit to a double Gaussian distribution.

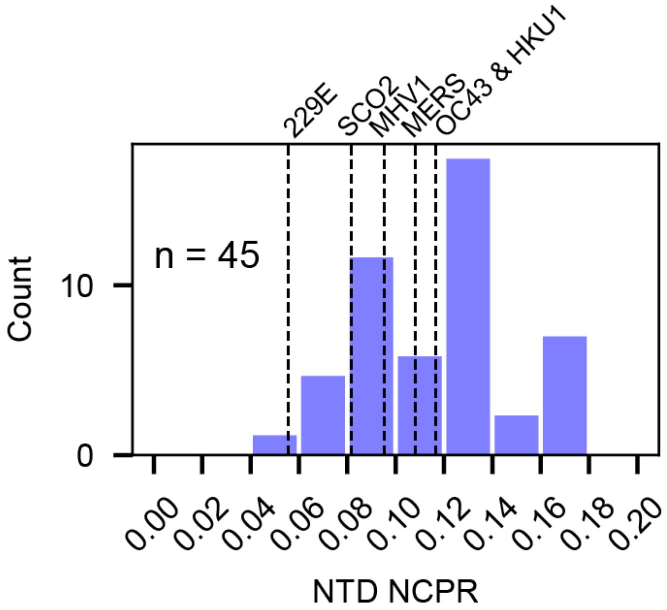


**Supplementary Figure 5.** Distribution of net-charge per residue (NCPR) for 45 different coronavirus N-terminal IDRs.

### Supplementary Tables

| Homolog | NTD Sequence |
|---|---|
| SCO2 | MSDNGPQNQR NAPRITFGGP SDSTGSNQNG ERSGARSKQR RPQGLPNNT |
| MERS | MASPAAPRAV SFADNNDITN TNLSRGRGRN PKPRAAP |
| MHV1 | MSFVPGQENA GGRSSSVNRA GNGILKKTTW ADQTERGPNN QNRGRRNQPK QTATTQPNSG SVV |
| OC43 | MSFTPGKQSS SRASSGNRSG NGILKWADQS DQVRNVQTRG RRAQPKQTAT SQQPSGGNVV |
| HKU1 | MSYTPGHYAG SRSSSGNRSG ILKKTSWADQ SERNYQTFNR GRKTQPKFTV STQPQGNTIP |
| 229E | MATVKWADAS EPQRGRQG |

**Supplementary Table 1. Coronavirus orthologs NTD**

| Homolog | Sequence (NTD and RBD separated out) |
| --- | --- |
| SCO2 | MSDNGPQNQR NAPRITFGGP SDSTGSNQNG ERSGARSKQR<br>RPQGLPNNT<br><br>ASWFTALTQHGKEDLKFPRGQGVPINTNSSPDDQIGYYRRATRRIRGGDGKM<br>KDLSPRWYFYYLGTGPEAGLPYGANKDGIIWVATEGALNTPKDHIGTRNPAN<br>NAAIVLQLPQGTTLPKGFYA |
| MERS | MASPAAPRAV SFADNNDITN TNLSRGRGRN PKPRAAP<br><br>NNTVSWYTGLTQHGKVPLTFPPGQGVPLNANSTPAQNAGYWRRQDRKINTGN<br>GIKQLAPRWYFYYTGTGPEAALPFRAVKDGIVWVHEDGATDAPSTFGTRNPN<br>NDSAIVTQFAPGTKLPKNFHIE |
| MHV1 | MSFVPGQENA GGRSSSVNRA GNGILKKTTW ADQTERGPNN<br>QNRGRRNQPK QTATTQPNSG SVV<br><br>PHYSWFSGITQFQKGKEFQFAEGQGVPIANGIPASEQKGYWYRHNRRSFKTP<br>DGQQKQLLPRWYFYYLGTGPHAGASYGDSIEGVFWVANSQADTNTRSDIVER<br>DPSSHEAIPTRFAPGTVLPQGFYVEGS |
| OC43 | MSFTPGKQSS SRASSGNRSG NGILKWADQS DQVRNVQTRG<br>RRAQPKQTAT SQQPSGGNVV<br><br>PYYSWFSGITQFQKGKEFEFVEGQGPPIAPGVPATEAKGYWYRHNRGSFKTA<br>DGNQRQLLPRWYFYYLGTGPHAKDQYGTDIDGVYWVASNQADVNTPADIVDR<br>DPSSDEAIPTRFPPGTVLPQGYYIEGS |
| NL63 | MASVNWADDR AARKKFPPP<br><br>SFYMPLLVSSDKAPYRVIPRNLVPIGKGNKDEQIGYWNVQERWRMRRGQRVD<br>LPPKVHFYYLGTGPHKDLKFRQRSDGVVWVAKEGAKTVNTSLGNRKRNQKPL<br>EPKFSIALPPELSVVEF |
| HKU1 | MSYTPGHYAG SRSSSGNRSG ILKKTSWADQ SERNYQTFNR<br>GRKTQPKFTVSTQPQGNTIP<br><br>HYSWFSGITQFQKGRDFKFSDGQGVPIAFGVPPSEAKGYWYRHSRRSFKTAD<br>GQQKQLLPRWYFYYLGTGPYANASYGESLEGVFWVANHQADTSTPSDVSSRD<br>PTTQEAIPTRFPPGTILPQGYYVEGS |
| 229E | MATVKWADAS EPQRGRQG<br><br>RIPYSLYSPLLVDSEQPWKVIPRNLVPINKKDKNKLIGYWNVQKRFRTRKGK |

| | |
|---|---|
| | RVDLSPKLHFYYLGTGPHKDAKFRERVEGVVWVAVDGAKTEPTGYGVRRKNS<br>EPEIPHFNQKLPNGVTVVEEP |

**Supplementary Table 2 Full Length Sequence of NTD-RBDs from each ortholog**

| Ortholog | Kappa | |
|---|---|---|
| | NTD | RBD |
| SC2 | 0.364 | 0.191 |
| MERS | 0.448 | 0.204 |
| MHV1 | 0.287 | 0.185 |
| OC43 | 0.254 | 0.213 |
| HKU1 | 0.278 | 0.191 |
| 229E | 0.392 | 0.206 |

**Supplementary Table 3  NTD-RBD orthologs Kappa values**

| Ortholog | Fraction Charged Residues | | |
|---|---|---|---|
| | NTD | RBD | Solvent Accessible RBD |
| SCO2 | 0.204 | 0.202 | 0.242 |
| MERS | 0.216 | 0.204 | 0.198 |
| MHV1 | 0.190 | 0.185 | 0.245 |
| OC43 | 0.183 | 0.213 | 0.263 |
| HKU1 | 0.183 | 0.177 | 0.237 |
| 229E | 0.278 | 0.296 | 0.425 |

**Supplementary Table 4. NTD-RBD orthologs fraction charged residues. Solvent accesible RBD are calculated from the average of their 5 AlphaFold2 generated structures**

| Ortholog | Net Charge Per Residue | | |
|---|---|---|---|
| | NTD | RBD | Solvent Accessible RBD |
| SCO2 | 0.082 | 0.040 | 0.066 |
| MERS | 0.108 | 0.032 | 0.052 |
| MHV1 | 0.095 | 0.000 | -0.011 |
| OC43 | 0.117 | -0.031 | -.053 |
| HKU1 | 0.117 | 0.008 | 0.0 |
| 229E | 0.056 | 0.088 | 0.126 |

**Supplementary Table 5. NTD-RBD orthologs net charge per residue**

| Ortholog | Hydropathy | |
|---|---|---|
| | NTD | RBD |
| SCO2 | 2.776 | 3.856 |
| MERS | 3.589 | 3.841 |
| MHV1 | 3.219 | 3.769 |
| OC43 | 3.333 | 3.854 |
| HKU1 | 3.210 | 3.790 |
| 229E | 3.378 | 3.713 |

**Supplementary Table 6. NTD-RBD orthologs hydropathy**

| Ortholog | Fraction of Disorder Promoting Residues | |
|---|---|---|
| | NTD | RBD |
| SCO2 | 0.776 | 0.669 |
| MERS | 0.730 | 0.651 |
| MHV1 | 0.730 | 0.687 |
| OC43 | 0.737 | 0.664 |
| HKU1 | 0.750 | 0.692 |
| 229E | 0.833 | 0.616 |

**Supplementary Table 6. NTD-RBD orthologs fraction of disorder promoter residues**

**Supplementary Table 7. List of scrambled sequences and their binding affinities**

Rep = Scramble #

| Rep | Sequences | Ka* average | Ka* std |
|---|---|---|---|
| **0** | SRGTSQGPNPDQKPIQQGSSGFNENRDPSMGTRAQGANNPRSNRRGNLTA | 2.363 | 1.540 |
| **1** | TRGGNLRQNNGTQSFAENTPPQIPDSGGGRNPKQDMGSSSAPRRNQSRNA | 0.922 | 0.248 |
| **2** | RNPNGGTQGRDANPRMGSRPPNTSSTLNSQQQINASNDPSKQGGFGRREA | 0.036 | 0.010 |
| **3** | SNSTFPNGAQGNGDNISRRSEARLQDGKMNSPQNQPPGPRQNTTGSRRGA | 2.417 | 0.351 |
| **4** | PNGSGNQNNNRSTLGSNSSGPRGPRMGTRSQATGAQRQDDNIEFPRKQPA | 0.032 | 0.005 |
| **5** | SEQANNSPTGGPPPPSRTRISQSNNMRQNDFNGKNTQGSGRRDQGLRAGA | 0.592 | 0.387 |
| **6** | SNGRGNAMTPNDSNQRRGNTFGLTSNKPPDARIGQGRGPPQSSQRNSEQA | 0.066 | 0.025 |
| **7** | GSDPTPGNKNNSAGNLSQGQSETGRIFNQSQNGRQDPMRARRGRPTPNSA | 126.308 | 88.768 |
| **8** | PQNLDSINFPADSSQNGNQRQPGTSSGTTMPRGNGGPGNRQAKESRRRNA | 445.878 | 144.100 |
| **9** | MNPTNGRRIRPDGNSNPFKASGQRSTNQDRSPGSGNEPSGQNAQTLQRGA | 0.099 | 0.025 |
| **10** | GPSGSGAGNNPSSQQRRSTQGRPNRDFTTMNNANQQIRRPGEPSLDNGKA | 0.074 | 0.029 |
| **11** | SPRPRQSNGGRGTNQMLNNQEPGISTGGQDSTANKFNPSRNRPGRQDASA | 0.085 | 0.032 |
| **12** | GRSNLMTAIKQSGGGNRTPPRQRGNANPSDQQESQTSFDGGSNRPNRNPA | 0.133 | 0.062 |
| **13** | QSTRTDFGQPRKDNQPRTRNPIGAQNSPGPRSNGSGSSNQNGENLARMGA | 0.023 | 0.012 |
| **14** | SSQQGSQPDTPRMIQRNEAPNRNRGSGNRPGSGAFTNQNRNGKLTPDGSA | 0.033 | 0.016 |

| | | | |
|---|---|---|---|
| **15** | NQQNRDTGIRGPKLNSPNNQNFRARSDRTMQEGPS NSPGPTASSGRQGGA | 0.096 | 0.042 |
| **16** | SGSTDGQPRRIGFAGTLPPSSGSNANSNQRNMKGNN RNDGTPREPQQRQA | 0.137 | 0.029 |
| **17** | NNRNGPPSQRQGQMRTPANNGSNSQNTPTQDPGSG RSEIRADFSKLRGGA | 0.097 | 0.017 |
| **18** | PMNSGNSRKPNNLNTGEFQSIPQPPTNQTADRGQD GGRGQRNSSGSARRA | 0.457 | 0.420 |
| **19** | GGNRRIPAKTTSSANGSNGNFPQSTQMSNQQGRDG RQRPLNRSPNGEPDA | 0.097 | 0.049 |
| **20** | QNRTGFNRMATGPPSRPSQSGASNILNRSSGDQEDQ NNKGQGPGPRNRTA | 1.438 | 0.929 |
| **21** | FTAGPPSNSTQNSKTNRRPSGGGDLEAQNMQPGDSQ RGNQRGPNRRNISA | 3.965 | 2.316 |
| **22** | PINTGRSPSQRGNNTDSQNPNQSNAFRGRPGKPDNA MLSSRTGGQRGEQA | 0.063 | 0.011 |
| **23** | QDTGPRSSNSPPSQNADANLTNRGGQTRRPGINRSK NQPGRGMFGNEQSA | 0.482 | 0.121 |
| **24** | QNSGEPGGDRTQRNDTGPSPRMFSRQQGRIQTSKAS PGNLARPNNSNGNA | 0.061 | 0.010 |
| **25** | QDPPNPQMNKNASNEADSPNRGSTGGGRFSGRNTQ RNQGSGILRTSQPRA | 1.995 | 1.549 |
| **26** | ERDTQDGNSFSRTQTLNPGSPNMPRNQSGPNKRGN GRAQSASGPGQNRIA | 0.528 | 0.433 |
| **27** | STRRGSDAPIQSKLSGPQRNSGDNTNPNANPSQQGP GQFRNRETNGGMRA | 0.053 | 0.012 |
| **28** | PSQPQRTQMNRGIDSTPDNPNSRNNGSQQRGPGSEA GRGTKGSAFRLNNA | 0.113 | 0.023 |
| **29** | GTMRQKDPSGTSTPGPNAGNSNRFNGRSLRNGDGR QPPQSNSNEARIQQA | 0.028 | 0.010 |
| **30** | GQGTGNNSQKRNSRMINSQTASPRGPGSQPRNNPLD TPFRDEQAGRGSNA | 0.068 | 0.027 |
| **31** | QNPARSQFGNRIRNTLRDSGNRGEAGDNPKPQGNP RSSMTNSGTGQQPSA | 0.033 | 0.015 |

| | | | |
|---|---|---|---|
| **32** | QIPNRTDSRKTPEGLRMRQGNQANGNSSSGTQPNSF APDGPNSRGQNGRA | 0.065 | 0.006 |
| **33** | NTRQSGGFNMRGTAETRSRPQANNDGNPPRPRQQQ SSSSGKGNDNLPGIA | 0.019 | 0.003 |
| **34** | GNNARIQRNPLFRERNNQTSDPPSSRQQTRGTSGSG QDAGGKMPNPGSNA | 0.027 | 0.017 |
| **35** | NRSNFPNPQSQLGQGAPRRPDGMNSRIAGNETDTSS TNSRKGNRQQGGPA | 0.493 | 0.289 |
| **36** | NPLNMDRQPSSTNNDNRQQSTRGGGIRGGSKPGRQ ARSNENSAPGPTFQA | 0.132 | 0.049 |
| **37** | NGGNRQNDNRQPITNFRNRRSPTPSKQASRQSPTGS NGQEGGLDMPASGA | 0.081 | 0.009 |
| **38** | KPTPNRLQPGRGPSSTNGRTDSQNSGNGAQPRGAFR GNQENQRSSMNIDA | 0.026 | 0.005 |
| **39** | QSIPNPNTLRNRPFRQSMDGTQSGRNSGTGAGRGNG DEPANQRQNPKSSA | 0.069 | 0.022 |
| **40** | NRIQQTGSPRQDMGQANNGQTPASGPNRTGRGRKG SESLPNPNRSNSDFA | 0.027 | 0.018 |
| **41** | SGRSSSSNRLQPGREDDGTGRPNPPNGQNNANRKQ TNPIQQMFTARGSGA | 0.188 | 0.116 |
| **42** | RKGQFRPGIQGPSSASDNNNTAMDNLQSERSSPRPN NPNGTGQRTGRQGA | 0.220 | 0.233 |
| **43** | QSAIANTTQGSSPSNPRRRQQGNRSDLRKRNGFSTNP QPGGGEPGNNMDA | 0.104 | 0.056 |
| **44** | SNLNGGQGPTGPIGRAMRPFAGNDSSTRNDQPRNQ RETKSPGSNQNSQRA | 0.078 | 0.010 |
| **45** | RSPPNQPDRNGTAGGTNSAPRDFQNNGSSNSNRIGQ MGRQKPLGRQETSA | 0.395 | 0.233 |
| **46** | SGRGQPPSSSMARSQNGRIRNGTPQDDPNRQEGFNQ ANLTGRTKNSPGNA | 0.041 | 0.010 |
| **47** | GNSGSDQANIPKAGNFNPPSQMQGSQGPTNGSLRER GDRPSTTNRNRQRA | 57.134 | 35.759 |
| **48** | GSGSKDQSGNQPFTSRGNGAGDANRNTLPNNPTQM PPRGSNQQERISRRA | 0.156 | 0.037 |

| | | | |
|---|---|---|---|
| **49** | GPQSGRNNIGASRANNDNFSQPGPESQRPNMTPSRSQRNTGGQGLDRTKA | 0.053 | 0.017 |
| **50** | FSSRRRNGPGRSGNRRSNNTDKSIATPGPQTAGSMQEQLQQDPGPNNGNA | 0.095 | 0.024 |
| **51** | GNQRAPRTRDQDGPRRQGNSQSSFQRSNNAIPNGNLGPSGMNGEKSTPTA | 0.057 | 0.012 |
| **52** | SMTGARAENGKNQFTGSPRQSQGQNTRRRRLPGQSPDPISNPGNNNGDSA | 0.138 | 0.081 |
| **53** | QMERSLSQRQNGTSRPNNQGAGGRRNGSDAQTSTRNSPPGDFNPGIPKNA | 0.302 | 0.139 |
| **54** | SNTNNGSRAQGNRTRGNQQRGDRRPISFPKGEPPMPANSGSQQGSNTLDA | 0.046 | 0.008 |
| **55** | RRRPGSGNGGGDQQTSRRPNRNSATADQNKQMSGLITFSNQGNEPNPPSA | 0.066 | 0.011 |
| **56** | SPNRSGDQQMGRNTPAPGNERPRSRGGNPQNTISFQLQNRANKDTSGGSA | 0.120 | 0.072 |
| **57** | QPNNQSNGNPRTSPARRQNERQGKGRRGQDSSPGSSIFMLANGDNTPTGA | 0.024 | 0.008 |
| **58** | RFTGNTLPDEGNRASQNNNSINRRGRTQDQGGPAQQPPRGKSGSSNPMSA | 0.101 | 0.024 |
| **59** | SQANRNDRSQTSRLESNQGRKANGGGRPRPSNNGSFGPMIDQPTPTQNGA | 0.042 | 0.014 |
| **60** | GRNSSNNTGNGQRKSIRDRTGNGSQRPREPDAQTGMPSSNFLAPGQNQPA | 0.025 | 0.003 |
| **61** | TRRRQQGNNGIRSQPMSGPARPPNSKLRSTNDGGQGNAFGTSPQNDSENA | 0.063 | 0.012 |
| **62** | RNRNSGPSGTGDSRRPKNGQNQRGSARITSPQDSNPLQGNPQFMETNGAA | 0.033 | 0.012 |
| **63** | NPESSGAISLRSARFNQRRDDQRSRGKTNQNPGSPTMQNNGQPTNGPGGA | 0.070 | 0.024 |
| **64** | NNRMPGSPNGARERRKPNDRGTGGRQNGSNSSSSNPQTIQTGQAQPDLFA | 0.033 | 0.009 |
| **65** | NARPQGQRTRNSTNRSPFQSNGGTNMARERDGPSGISSDQNLKPNGGQPA | 0.059 | 0.020 |

| | | | |
|---|---|---|---|
| **66** | NNQRISRSGGRKNGETLAMSQRSRQAFRTGPNQQTS PPGPGSDNNGPNDA | 0.018 | 0.004 |
| **67** | FRQLSIGSAMDSRGPESNSQPNARRTRKRGDNGPPTQ SNNPNGGQTQGNA | 0.089 | 0.027 |
| **68** | GGRRFSDPAQPGPKTSRQTGNNRMLRRNSNEIANQD STPGNPGSNGSQQA | 0.131 | 0.065 |
| **69** | QGGRSNSPSRIKTDNRGMNGLENTDRRGRPQSASNP QNTQAQPGPGSNFA | 0.043 | 0.012 |
| **70** | TPSTQMKRPPIASNNESRNRNSGRSRQDRNSQQGQG DPLANGGFPNGGTA | 0.081 | 0.024 |
| **71** | NRPNKGNEDSPGTDQQPTRRRPRSGSRIQGNSAQPS QGNTNGNMSAGLFA | 0.111 | 0.049 |
| **72** | NGPNPNGPLDQRIGKMRSSQGRESGQRRTRSAGQD NFQSTTGPANPNNSA | 0.078 | 0.017 |
| **73** | RQKRSDDQGNGEGRGPNTSQRRGPMFNGQRSATNP TLSGNPANNPSSIQA | 0.032 | 0.011 |
| **74** | DRRGKSGPPPTRDSQMNTANRRTSPFESLRQNQQNS QNSAGPGINNGGGA | 0.025 | 0.007 |
| **75** | RNGSSPRQNGRMNPNGQDKRNQSRPIPSQRSNAQN GGDGGATFESTLPTA | 0.030 | 0.014 |
| **76** | QGTLKTNNRRNDNRQPPMGRGGNAPRNGPRQNQD TIGSGFEASSSQPSSA | 0.026 | 0.005 |
| **77** | RNNRREQRQGSTTKQSNSSGDSGRPRPFAPNGNLQS GGDNQPNMAGPTIA | 0.021 | 0.007 |
| **78** | STKNRRQMGIPPSNRENDPRQQANGRPRNSGGTTSD GLGFNQSPNQSGAA | 0.024 | 0.022 |
| **79** | RGDRAPNRSSLRGPPNSQSQQRSRKQEFTATNPGTQ GGDGGNPNNMISNA | 0.066 | 0.015 |
| **80** | PPRGMDTSSSQPGNRPNGRTRRQRAQEGKDTNSNAS QPINFNNGSGGLQA | 0.052 | 0.010 |
| **81** | GRQREDIQQQATNPPPGFSTGKRRSRAQSRNTGGNN PLSMNNNSPGSDGA | 0.528 | 0.128 |
| **82** | PRRMDGTQGRPRLGGPPQNNKSNANSRQRSENQGS STGDNQTPIANGFSA | 0.031 | 0.009 |

| | | | |
|---|---|---|---|
| **83** | SSERQMQIGPRNNGNGSARGTKRQGGSRNRPFDNT TGQLDPAQPSPNSNA | 0.037 | 0.007 |
| **84** | TEQPFQTGQKRNPQNIGSRRGRTNGRRPDAMNGNS SSDAPPSNQSNLGGA | 0.212 | 0.046 |
| **85** | NGKRPGTGSNSRQNTSPRSQNSRATRSRGPIDQMEG QNGQNPLDAPFGNA | 0.047 | 0.006 |
| **86** | RPRQASSGIRGKNFGTTRGNQDRGNQQGSRPSTQAP GNMLSEDNPNPSNA | 0.025 | 0.004 |
| **87** | DFAPTDNSQEGPMSQRGRGPKARRRLGRPSTNPSSTQ GNSQGNNNIGQNA | 7.102 | 2.195 |
| **88** | QGRRRANMTGNIQQNRQPRGNREGANPKLTSSPQD FSTSGNSPDNSPGGA | 0.054 | 0.006 |
| **89** | SKGGRNPMNQPDNNLQPPTRRRSTENARGRQGFSG QQSTNGASDSNPIGA | 0.030 | 0.005 |
| **90** | RRRSSGTGPNQNQLRPGQKNGGPSRQRDDTITANP MGPQSNSNAGENSFA | 0.039 | 0.005 |
| **92** | NRQQNNSQETDRAGFSRSGMGLRTSTIPNPSNGQPA DGSPGRKPNGRNQA | 0.136 | 0.044 |
| **93** | GNNENQDPQTSRSFTNKNSRIDNSPGGGAAGQNLP QPRTGGMSQPSRRRA | 12.046 | 8.509 |
| **94** | PPPARSKRDTGGNQRNGLSQTDRGMARENPRQNGQ PSTGGSNSQNFNSIA | 0.018 | 0.006 |
| **95** | EGQRSPGTSQTTGDNNRQSNSGRQSARRNQSFMPAD IGPNKGRGPNPLNA | 0.103 | 0.029 |
| **96** | PFGLNNQPGNGRPMGRRGANEPGPKSQSNNSQSTT DSSIRGQNATRRDQA | 0.019 | 0.017 |
| **97** | TTTRNQPNRIGQPNNPKPQNQNARSMEGGRSQSND GSAGRGSDSFLGPRA | 0.019 | 0.004 |
| **98** | NDDNSTPGPGPRENFTGSTNIRNQKRGGNQSQGAP RQRGPANSSQLMRSA | 0.368 | 0.097 |
| **99** | STGEKRTSSGLIGDQGQSNGQGPMRPNGNNSQATP QRANNNPDPRSFRRA | 1.784 | 0.889 |
| **100** | SPGQSSQTNKSRGGNGLGTNDPNPGFTARRRSRPRQ PEDQNAGMNSINQA | 0.339 | 0.099 |

| | | | |
|---|---|---|---|
| **101** | DGIFQGSNPGRGLQPSQGNGESNDAQNTKSSTARPQ TGMSNPRRNNRPRA | 40.181 | 27.957 |
| **102** | NRPLDTRPSQGGNFQSQIRAPENNGRGGNRRNTMK PSGGSNQATPSSDQA | 0.057 | 0.017 |
| **103** | GTDQSNGTRQPEQSNPNIGNRGRLRNNPSGNQRASF PGMGKSDPTSRQAA | 0.658 | 0.269 |
| **104** | RGPQGSTFDNLRSSGTQQNRPNMGIRPPKRNQQNPR GSDGSNSANTEGAA | 0.070 | 0.015 |
| **105** | KRNFSANTDGNGTQDRGSQNQNGPAGTPMESNQR PLSPSGQIPRRGNSRA | 0.784 | 0.259 |
| **106** | NRRFANNRSGATKRSQQQSPTNTPEDSSLQQPGNIPP GGMGSGDNNGRRA | 0.084 | 0.025 |
| **107** | RGAGSNMATQTGNSRSGGQPPQKETRQPLNSQFNN RGNNSRPSRDIDPGA | 0.039 | 0.013 |
| **108** | GGTKNSNSRRSQDNATRTASRNPPNDGPIPPSNSGQL MREGQQFQNGRGA | 0.044 | 0.009 |
| **109** | GNRGRASRPRNSTNFLPKRMSTDPQTDPGAPNQGSG NNGRQQSSQEIGNA | 0.072 | 0.033 |
| **110** | STGSGPRDIQSKLTRRQTMRSDRNNQNANQGGNPN NGFERQPAGPPGSSA | 0.052 | 0.006 |
| **111** | SRRGGNPRPGGQSTSNPTGNRNRERSGQTIDSDQPF QANQPMLSNKGANA | 0.026 | 0.010 |
| **112** | AGPSGSNGTQSPNDNKQGRNRPSQDAQGILEPRPST GNRFRSTMGRQNNA | 0.562 | 0.125 |
| **113** | NRNSIDNNRQRNPSMTERGQATQTSGGPRKNGSGG PGASRPQPLNQSFDA | 0.038 | 0.009 |
| **114** | SLAERFQITQAGNGRRRPGQPGSMNTSQPGNNNGRS TPGRNKNDQSSPDA | 0.078 | 0.009 |
| **115** | GNRPMGLNQSSGTAINQKGSRNTDFDGNQSSPGQR QPTGANRRPPESNRA | 0.148 | 0.045 |
| **116** | PPQAGAPQQSTMGSRRFNQINPNETGTKNQDPGSRR DNRGRNGGSSLSNA | 0.058 | 0.008 |
| **117** | TNQESPFRRGNSGPPLRDTQARPQGKQGARNPSIGN GMTGRNSSSNDNQA | 0.044 | 0.019 |

| | | | |
|---|---|---|---|
| **118** | SPNQQARSNNNTPPFPERGTIQQGRGLRMSGSNSPD NTGKGSDRGRQNAA | 0.037 | 0.011 |
| **119** | STNGPSSNRQGSGDRPNGGFPLNTQGNDGRSISQAQ MAQKRPEPTRNNRA | 0.126 | 0.050 |
| **120** | SKNGEQQGRRGNPGSDMAPFRINQRDTGRTPNQPN NTNSRQPALGSSGSA | 0.028 | 0.007 |
| **121** | SQTQRNSGSKARRNRERRPFGGNNMTNNSQITPLGP AGPPSNQDGDSQGA | 0.099 | 0.026 |
| **122** | QGTTSRRARRDRGNKRAGGGGQPTSSNNNPNSPQE PSSNFGIMNLQPQDA | 0.232 | 0.049 |
| **123** | MSAPRNRRERFPSDKRARGNQNSSTSGTNGNSNQQP QTDPNGGPLGIQGA | 0.060 | 0.008 |
| **124** | SNEPRQRGRRSKQPARNRMQSGNGSNQDNDTNSPSP GTGIAQLPGNFTGA | 0.154 | 0.067 |
| **125** | GGAQRRRRDGQNRRFSGKIGNQNQNGSPPALNSGM QTNPPSNTESTSPDA | 0.131 | 0.081 |
| **126** | SEGNFRRQMRKSDRRRNQPGNPSSSDPNTIQNGSQN TLGQNGATPGPAGA | 0.278 | 0.127 |
| **127** | LNNERRQTRPRPPRKGGRGFSGQSIMNNNTTGQSPQ SDGADPGSNASNQA | 0.094 | 0.048 |
| **128** | DNTSPRPQRNGRRNKRRPLGTAEQQSGPFNMNGND GIGSSGQPTASNQSA | 0.393 | 0.161 |
| **129** | NFPSGGSGRQNRKSRRRRNSSNEMPNGTADQLPDQA INGGNGTPQSPTQA | 1.065 | 0.550 |
| **130** | GNFGSGRQRRRNKQRPARQPSPSTGDIGPTQDSGQL AGSNNNPESMNNTA | 0.194 | 0.030 |
| **131** | PSNQRDRRARRGNRGNKDQTSPNGFMATLGQPNSN EITGSQGSGPNQPSA | 0.077 | 0.037 |
| **132** | LGNDRRRPGRRGSNFRKNPQANQSTMPNDGTSTSE NNPIPSGAQGSQQGA | 0.200 | 0.048 |
| **133** | QLAQRSRRRNRNIKPGFRQMSGGNSEGDGDTTPQAT NGNQPPGNSSNPSA | 0.402 | 0.211 |
| **134** | ENQNSPRRQLKRRRRSIGPNGTSMNQQQADSTSNPP NGATGDGGNSPGFA | 0.889 | 0.201 |

| | | | |
|---|---|---|---|
| **135** | PSNDNSRRQGRTKTIRRRNGAFSANPGGPGPGNQPQ MSQTSGSLNEQNDA | 0.211 | 0.017 |
| **136** | AGQNTRRRSRNRNGKRSPTQQSMDFGNSDPEQPNTS SIAPLNNGGGQGPA | 0.246 | 0.033 |
| **137** | QADSSPGRANRRRRRKQQNTPIFLNGGSPTDNGSPQ ESQTNGPGMSGNNA | 0.645 | 0.342 |
| **138** | NNGPLRRDSQRSMRKPRRGPNPITNDTGGFGEAATN GQGQPQSNSSNQSA | 0.189 | 0.071 |
| **139** | TSGQENSRPNRRAQRRKRNMSSQGGLFPGPSTNNGD DQAPGSQTINPNGA | 0.638 | 0.171 |
| **140** | TQSSFRRMERKQNNGRRRGPDSQGQPINTNTPGGN GANPLPSSQSNGDAA | 0.099 | 0.029 |
| **141** | DSAAKLIQRPRNRRRRFTDENGSSNGQGGSPGQSTN NPGMNTSPGQNPQA | 0.698 | 0.331 |
| **142** | TSEGRFTRRNGRGRRPKTNGQQQAQNSNSGLMPQP NIPASNPGDGNDSSA | 0.252 | 0.079 |
| **143** | NENGGNPRLRPNKRRRRQSGQGMNPTQNDSATSSQ NGSFGSIGDAQPPTA | 0.286 | 0.220 |
| **144** | NFDERRMNRRARNKPRPSQNNLPTTGQNTDPGSGQ SNGSSQSGGAQPIGA | 0.115 | 0.034 |
| **145** | PAFNQPRGERRRSRRGKPASDDSNQNQNGGQMTSN LGNSPISQTGPTGNA | 0.271 | 0.158 |
| **146** | SDTQRANRRLRPTKRGMRQGPQPNPQDENTGISSNA NQGFSSPNNGGGSA | 0.379 | 0.202 |
| **147** | NENDRRAGGRRRGTGGRKNGQSSNSDNAIQTSPQP FPPNSMGTSQNQPLA | 0.160 | 0.038 |
| **148** | NQGNRRGGNTRGRNKRRDTSMINEDQSPPFQSSGAS ALQPPGQNGSPNTA | 0.050 | 0.042 |
| **149** | SSNQSGRRTRRLRRESKTAQNGGQINGPMNPDGFQN SAPGDPQPSTNNGA | 0.232 | 0.031 |
| **150** | NAGSGRPNKRTQRGRGRRFNQSTPPENISGSLNNTD GNDQSSMAPGQPQA | 0.360 | 0.052 |
| **151** | NNASQNKRRRSPRRGQSRPNNDTPGQADIGGTPLQS NMSQPSFTGGNEGA | 0.204 | 0.096 |

| | | | |
|---|---|---|---|
| **152** | TGDNKGPRRSRQRSTRRSGSMQNGGDIQNPPNTAGP LPNFSANSQQENGA | 0.196 | 0.067 |
| **153** | DAPQRSGPRGSKNRSRRRDNQGTNTQSGGNGAEPP TSLNSPNGNMFIQQA | 0.277 | 0.114 |
| **154** | QSPGRRRSTNRQDRRKGNSSTSAQGGALDNNFMPN GETGPNPIGPNSQQA | 0.109 | 0.017 |
| **155** | NPGSKRRQQFRIRNGDRRQPNGNGNPSSSEGLSGTN NADPMSGQPTATQA | 0.069 | 0.022 |
| **156** | GSNTGNDKNRRNRRTRREGIQSQGGFPNPMSLQPST NQPQGSSGPADNAA | 0.514 | 0.082 |
| **157** | QGTNSRRRGRRKAITFNRPSENPPSMGAPNNGNQSN QTGQSDGGPLDSQA | 0.156 | 0.051 |
| **158** | DGGQRKGSNNIRTRRLRRMPNTPNSPASGNDQPQSN ANSFPSGEGTQGQA | 0.201 | 0.174 |
| **159** | NPGNRNRRKQRRRGINESGTSFQPNALQSGPGPPAST QNSTMSDGQNDGA | 0.229 | 0.045 |
| **160** | SNFGGRQPSNRKRARARRNDNPQETSGPSQMGLQNI NNTTDPGSPGGQSA | 0.343 | 0.187 |
| **161** | QADTGRLTKGRGRPRTRRGNPSSNQNMPSEANQGIN SGPSSGNNFQDQPA | 0.272 | 0.136 |
| **162** | QPATQRRRRKQERSNRPNPNFAMTPQSSLPNGIQGT DGNGDGGSSSNNGA | 0.328 | 0.140 |
| **163** | QTSPGKNRNFRRPARQRRSPNPTSGEGQDQSGANLG QSTNNGDNSGIMPA | 0.245 | 0.033 |
| **164** | NEGAFSRRTPGKRGRRQRSSPPQNTNDTSNGNDGSM QPSGNQIQANGLPA | 0.418 | 0.140 |
| **165** | GQDPPRGKRNSNRRMTRRFNAQPGNTTSNGDLGQE GQPSANSINSQPGSA | 0.322 | 0.037 |
| **166** | NQNIGPRRSNKRRDPRRLPSTMNGQGGQEQGDQNS AGNSFSGNPTAPTSA | 0.249 | 0.119 |
| **167** | PNGSRRNERQPNKRRRTQPAPGDIGTNFPGATQGQ NNLDGQMSSSGSNSA | 0.096 | 0.024 |
| **168** | TNSTRELRQKRNRSRPARFPAQSGNGQGGNPGISQD GNPSSTDNQNPMGA | 0.130 | 0.034 |

| | | | |
|---|---|---|---|
| **169** | FQQTRNSTAKRRRNIRGRSNPGESGAQSSPNMSNQG PGGLTPGDQPNNDA | 0.242 | 0.035 |
| **170** | QANDRNFRRGGGMKSRRRPNSTPITLQATNNQNEQ PSSGPQDGNSGPSGA | 0.121 | 0.034 |
| **171** | ANNPNGNGEPFQQRRRSRRRGKSSTQDNPSTGNGG SSNAQTLIMGPQDPA | 0.992 | 0.385 |
| **172** | QDTQSNPPIGNQNFQKRRRRQRRSEGGGTASSGGA GLDMTPSPNNSNNPA | 1.904 | 0.655 |
| **173** | GPGNSSEQPFNGTRKRSRRSRRLAMGIDATNTPNNQ GQNSSQPDNQGPGA | 2.148 | 0.185 |
| **174** | GQASGGNPQPNGNKRPRRRRRGDISDPTGSSNQMA NQQPTNEFNGLSTSA | 0.892 | 0.311 |
| **175** | GTPITGSQSPNPMRRKRNRRNQRGASPGSANQLENG FQGDSPQDTGSNNA | 1.607 | 0.559 |
| **176** | NGSNSPIPLQDGGRKSRRRDRRQSNANNNPNFQAGT GGQSTPQMSGTPEA | 0.187 | 0.028 |
| **177** | DGAGPTGNPGNQMRQRSRSRRKRSNPNTGPFQNGT SQLDGQISEANPNSA | 0.654 | 0.073 |
| **178** | AQINDTPNNQSQGRPRPRKMRRRAQNDGNGPGTNS PNFSSESLGQTGGSA | 1.040 | 0.384 |
| **179** | DANPQGGSSANQFRKRRRRRQPTESTLTNGDMSPPG GNNSNSPQQGNGIA | 1.054 | 0.161 |
| **180** | QGQPQPPLFAGNQRRKRSRRGNRNGDMEGSQSNDS TGNSTNANTGISPPA | 1.327 | 0.151 |
| **181** | PNNQNDSSTPGDGRRFARRQKRRTENPGSNSSQLQM GGPNIGTASNQPGA | 5.569 | 3.532 |
| **182** | PTTQNGPSNQSLSRKRRPRRQRSNQDPIDNNAGGEN TGPNMQSGFGASGA | 2.442 | 1.277 |
| **183** | NTQFSGSQMGGNPRRKARRTNRRPAPPISGDQNQGP NLNGSQGSTNSDEA | 0.373 | 0.060 |
| **184** | TNNDPQTSSSGGARRRRKNRRGFANTQLGGPGMQP ESNPQSINPSGQNDA | 1.668 | 0.320 |
| **185** | SNSQGPNTQPNNPIQRKRRRRRGSTPSNAPNGSQGG MLGETDAFQDGNSA | 2.218 | 0.560 |

| | | | |
|---|---|---|---|
| **186** | GDNQSNSNGNPQTRRDRKSRGRRPTGSQPMASPGTEIQNLGFNNQPASGA | 0.770 | 0.265 |
| **187** | PPSSNEGTGQTNPRKFRRRRNRSQSLNAPANGPQNSSGNIQMDQTDGGGA | 0.838 | 0.124 |
| **188** | GAGSEGNIGDTLPRRRRQQRSKRNFMTAPSQGTQNNQPSNSNPDGSNGPA | 1.204 | 0.223 |
| **189** | PGIGMPGSNQPNQGDRRRRRTKRQASTNSGANQGSNESTDNLSGQPPNFA | 1.120 | 0.874 |
| **190** | GFISMSSDTGNGGRRNSKPRRRRANPQDNNPQTGNLAQESGQPGNTSQPA | 2.370 | 1.163 |
| **191** | IGFQGPALESTGDRRQRRQRKRQGSDNNGAPNSSSGPNQGSNTTMPNPNA | 0.105 | 0.077 |
| **192** | PQTPQSGDGSQTTRRGIRRKRPRAFNGSSNNPGNQGDSQMNLESNPNGAA | 2.097 | 1.167 |
| **193** | STFQNNDGISAGGPRKRRRRNQRGSQPQGLANSPPGDNSPETMQSGNTNA | 0.946 | 0.408 |
| **194** | AQEGQSGNPTSPGRGRNRKSRRRFGQLPDAPNNTSGSNPTSGNQNDIQMA | 2.731 | 1.086 |
| **195** | NQAMSGLPTAGFDRIRGSRRKRRDNSQNGTGPSSNPQPSNQPGNTGNEQA | 0.271 | 0.090 |
| **196** | SPPEPGGNSGQQGTRKPRRRNRRFIGNNDQGSSSSPAALTTQGQNNMDNA | 2.561 | 0.737 |
| **197** | SNINQQGMGNPNARRRPKQPRRRNTSETSGSDQGGTDGPPNGSQFALSNA | 0.265 | 0.093 |
| **198** | PNSTSQPNPTLSGGIRKRRRRRQNPFGNQDSNASSGTEQANGPNMQGDGA | 2.935 | 1.443 |
| **199** | NQNGMFNGTLNAPKRINRRRRRSTQQEDGSDTGSQSPGNPSAGNPPSGQA | 1.213 | 0.784 |
| **200** | TPMGPGGFSNNGIRRQNKRRDRRQTSEDNPLPSGQGAQSSGNPSQNATNA | 0.268 | 0.084 |
| **201** | PQQQIDQSLENTSRDRGRRPRKRMQTSGSFAPGPSNNPGNGASNGTNNGA | 0.768 | 0.291 |
| **202** | NGSLGPGPEPNPGNRRRTRQKRRTQSDNINGGGQMSSDNSQAQPNASTFA | 0.757 | 0.475 |

| | | | |
|---|---|---|---|
| **203** | NGDGSPGSFNNGSRRQRKRRIRGGDNSPQQGTPLQP PTAMQTSSNNNAEA | 0.481 | 0.377 |
| **204** | SNTGPGMQSTGDSKRRRRRRPGAGNSNAQGNNDQI NQSFPPPSQNGETLA | 1.291 | 0.568 |
| **205** | GQSQQGSGNLAINRSRRTGKRRRNFTGGNPAQPDPP PNSMTQNSESDGNA | 0.782 | 0.253 |
| **206** | APPQGQPGQLDGSRKTRRRGRRNNNGASSQMPFGN GQNSIDTENSSTPNA | 1.216 | 0.335 |
| **207** | PQNMGLGSQTSNDRQRNRKRRNRTQGIPPNAATNE GQFGSPDSGGPSNSA | 1.273 | 0.488 |
| **208** | GQQFNLDSNGQAPRRSRNKIRRRGPPNASDQMTSGG NTNPEGSNTQPSGA | 1.210 | 0.503 |
| **209** | NGPQTSANGPNAQGSRRRKRRQRELIGGQPNSGQN DNMTTPSNFDPGSSA | 1.831 | 0.897 |
| **210** | PGSSGSQQSPQQPKDRRRRTRNRFGQPEANDTGISL GNMAGGTNNPNNSA | 1.036 | 0.913 |
| **211** | NETQPNNSDPPPNRLFRRRKGRRGGNSNDSMGSQSQ GTAGGQQIATPSNA | 1.139 | 0.535 |
| **212** | QQQSFTSDPTPNSRRRKRRPNSRGPGPIGNNSDNGA GQATEGQMNLSNGA | 0.701 | 0.117 |
| **213** | SSGISQTPPGGNSRARRPRSRKRGNNQQTDLAQNGSP GNEGFPMNNQTDA | 0.310 | 0.112 |
| **214** | TGLSQGMPQGGPSRPRNRRRNKRSDTQANSTGSQG QNNADNSEGNPFPIA | 0.895 | 0.480 |
| **215** | QFMATNGGPSNGNNRRPRSRRKRTGNGSQEDPDIQ PSPNQTGGANQSSLA | 0.878 | 0.545 |
| **216** | GQGFNIEMGGQNDRPPRRNRRRKTNDSGPAQLQNQ NASPTTNSGSSPGSA | 0.950 | 0.797 |
| **217** | FGPAANSNNPQDQRKQRRERRRGGIPNGQPTSGPQL MSGDNTGNNSSTSA | 0.225 | 0.088 |
| **218** | GANQNEIQNGNGSPDRRRRRRKSTPPSLAPQNFDTQ SGTGNSQGPMNGSA | 4.596 | 1.743 |
| **219** | ALPSNMQNPADTGRDRRGRRRTKQNNSGINQPPSEQ GSTPSNGQGNSGFA | 0.769 | 0.437 |

| | | | |
|---|---|---|---|
| **220** | GDFPSANNLNQNSSRNKRGRRRRGQASPPPMQNTG NSQSTGPTEGDQGIA | 0.451 | 0.099 |
| **221** | SEGPNTQGINSSSGMPSARSRARRRKGRDNNTQGTPP GNQQPNGFNQDLA | 0.620 | 0.603 |
| **222** | PPFDNMTIQGGGGPNASTSRRRQRSRRKSNPQNNGD NQEASLTQPGSNGA | 9.138 | 2.387 |
| **223** | QSPGETSINAQQPMFGSNRRRRGDRSKRPNPGGANT GNQSQTDNPNGSLA | 1.428 | 1.199 |
| **224** | TPGAGNGNNENGNAGSSSRRKRQNRTRRPSFGPPSD PQLDNQMQGQISTA | 1.043 | 0.521 |
| **225** | SGTLFQPNNIGEMASSDTQRRRRRSRKQPAGPGQPD NGNNSNQTGNSPGA | 10.939 | 5.654 |
| **226** | MGAPLNNDEGNNSSNNSTRASRRRRKQRPQGGSQN PTPGQSTGPDQFGIA | 4.205 | 1.600 |
| **227** | SPGNPGTGISQQGANGTPKRRRRRNRPTPESGLQDG NFMQSQNNASSNDA | 3.707 | 1.265 |
| **228** | FSGAPQNSPSSIGQMGQNRPRQRKTRRRGPENNGDS NLQNDANPGGTTSA | 4.854 | 1.766 |
| **229** | GQSNSSNDGFNQSGGPDPRRRPRKRRSTQAGGIGSN NTAQNMNPPTQLEA | 2.171 | 0.765 |
| **230** | NEGPTSPSPGNAGMFGQPRRLRTKDRRRGATPQQIS GGDNNNQSQSNNSA | 1.083 | 0.747 |
| **231** | QGGPESQATQSNMPPSNGRRRRLRSPRKGNNNQNP DADGSGFNTGSTIQA | 2.122 | 1.629 |
| **232** | NQNPGFPGNSSPSISGQARRTRQRKRGRQPTPTDNQ AGNGLDENGSSNMA | 2.315 | 0.774 |
| **233** | FNAGGAGTQDIPSSNTQNGRRSKRRLRRPNSGNGMD NNQPGEPPSQTQSA | 1.786 | 1.214 |
| **234** | TPSGGEDNGINPDAQFPNRGNKRRRRRQASGMPSSQ NSPGNTGQSLQNTA | 3.996 | 3.993 |
| **235** | QIAESANSSQDSQSNTPGNRKRRGRRPRNFGTNGGP SPDMGQGNLPQNTA | 13.916 | 4.699 |
| **236** | MNQNPFESNISSSGTGNARRRRPRPRQKPGNTALND GNDSQGGPGQSTQA | 3.803 | 2.379 |

| | | | |
|---|---|---|---|
| **237** | PDDIGSGMTSQPNQTQNNRSKRRGRRRSAQNGGNP TEFPSANPLGNSGQA | 1.406 | 2.077 |
| **238** | MQISPTSQNGQGNLPQPESNRRKNRRRRGPGQSFNT PGGTAGDSDASNNA | 1.294 | 0.324 |
| **239** | LNGGGPNNSNSAPSSPSQRRRRRDQIRKNGAPTQFN TSEGNGDGTPQMQA | 2.200 | 0.907 |
| **240** | PDANAQISMTFSNNNNPLRKGRRRQRRNGSTESGNG GPGSGSQQQDPPTA | 0.669 | 0.298 |
| **241** | PGSENNGTGQGNLPQQSSRRRRKSRFMRGGTAIGSP NDANQDPNPTSQNA | 1.671 | 0.310 |
| **242** | GPNNPGQPADSGNQGNPANTLRRKRRRREGNSQQF DSNSSTPQIMGSTGA | 24.669 | 9.803 |
| **243** | QTNPPDGEMNANGLNFNGRRIQQRRRRKSQTTPAQ SDSPPGNGSGNSSGA | 1.205 | 0.530 |
| **244** | NNQPGDQGNASPTMNGTIRSRKRRRDNRSGPGSLQ GSAPQNTSEQPNFGA | 1.012 | 0.937 |
| **245** | MGQSEIGGPLQNQNNAQSQFTRRRRRKRTSNSNTG NSDAGPPNPDPSGGA | 1.388 | 0.339 |
| **246** | NTNNEGDSSDGNSPGQPNRRQKRRRTRGFPPQQNIA GSQSSMPGNAGLTA | 6.060 | 1.443 |
| **247** | GPTFNSASPDPNGGPDSQKRRRSTRRGRTGGMNSNN ISPNAQLQEQQGNA | 1.341 | 0.413 |
| **248** | LPSAQQNGGTTNGGPSPQRRRRKRIRFDPNGDPGNT MNSNQAGESQSNSA | 0.709 | 0.327 |
| **249** | NPSNSTQTTGLPGPDQAPRKRRGGRRREQPGNSMNI FDQSASSGNNGNQA | 3.881 | 2.487 |
| **250** | SNSSGNGGTPTADGNGEPRQRRRITRKRPNSDPNGQ AQMFNLQPGQSNSA | 1.366 | 0.469 |
| **251** | STGAGIPNQQEQQPGFPDGRRRRRRDTKSANSNGLM SNGQSPGPNTNSNA | 3.216 | 2.039 |
| **252** | STATSNSGNGQNGQPQGARRPTRRNKRRSSQDEFDG INNMGGQPLNSPPA | 1.760 | 0.617 |
| **253** | GASSNFAGDNGQENSPISRRRKRNNRDRQGPSNLST NMGQPQTTGGPPQA | 0.701 | 0.306 |

| | | | |
|---|---|---|---|
| **254** | SQMQPGPTLDPENGIDFQRRARRSRKTRGSGGSNNSNGAQQPNPGSTNNA | 3.154 | 1.344 |
| **255** | IQSEMGGNQSDQPGFTGTRRGKRRQRDRSNPGANGSSSQPNPLNTNAPNA | 0.390 | 0.104 |
| **256** | NNESGGANTQTDQQSLSDRKRSPRPRRRPNGQMAGSNSQGFGGINTNPPA | 3.684 | 1.473 |
| **257** | EQPGDPDLGNIFATQGNSKRRRRRPSSRNASGSGPTNPGNTQGSNMQQNA | 1.097 | 0.714 |
| **258** | PPGGPSNSSSQADQFNNGRNRARQRRKRNGNLETNPQSQSIGTGPMGDTA | 0.748 | 0.289 |
| **259** | DNTNNDGPTGSSASNMQNRRPRKRRRQGLEQIGGQNQAFPGPNPSSTSGA | 6.686 | 3.235 |
| **260** | NPGNGNGPPASEMTPGQDGPRRRRKRSRTIFLNQGNSGQSQTNNADQSSA | 13.981 | 3.187 |
| **261** | QNPGAGSFPSDQSQPNNSQRGRRRRRKGNTTMEDNAGLPNTQSGGNSPIA | 7.407 | 2.099 |
| **262** | PANNGFGIGQTESDSPGQRRKPPRRDRRNQTQLTSSAPQNNSMNGSGGNA | 0.149 | 0.054 |
| **263** | NDQQPNANSIDNSGEGGPRKPRRTRRSRASSQGQPGMFQPNSGTTGNLNA | 7.696 | 8.015 |
| **264** | TGPQQQGEANISPLGSSDQRKRNGRRRRGTSNNPNNPSMGSFGPNATQDA | 8.431 | 1.907 |
| **265** | NGQGSPGSDPSNMASQNGRGPRRRRRKDNTQTSEPGTAQFPILSNNNQGA | 4.790 | 2.640 |
| **266** | NGGQDMGAPNQSGNSPPIPRGRRRQRKRSAPLGQTTDGSETSNNNFQNSA | 3.219 | 1.561 |
| **267** | QAPGTGDNNPQPNGNQPDKRRRRSSRRFGSQITAGNENNGPMLSTGSSQA | 3.725 | 1.624 |
| **268** | TAGGEQNSDGPLQSPNNTNRRRKGRGRRQSPNSADSSNQFQNPPGTGMIA | 9.709 | 5.751 |
| **269** | ANDNSPQIGQSPTPGNSTRQRSRGRRKRGNSDPGPNASTNQMGNEFGLQA | 6.552 | 1.044 |
| **270** | GNNQGGNMSQTQGSGTNFRDRLRRRTRKSPPSPNIPNSPAASDQEQGNGA | 0.751 | 0.368 |

| | | | |
|---|---|---|---|
| **271** | TQGGGTQNNSDNAGNGQDSPGQFKTRRRRRPNRIS MSNQPAPSESLNPGA | 28.752 | 12.245 |
| **272** | GQQEGSNSPNPDNTLGMNINTPSKRNGRRRRGRGG FSPSSADQPNQQATA | 78.551 | 63.623 |
| **273** | FTQQNNGSSGAINGQSDTQPTSERRKRRARGGRSLPS DPNNGGNPMPQNA | 0.787 | 0.454 |
| **274** | GSNAFAGGDTMTTQGGQNQNNGPRKRSRERRRPSN QPLNSSDGNPSIPQA | 0.535 | 0.166 |
| **275** | QNESMTQGPNGPNNDDQNSGNLFRRRSRRPRSKSSG AAIGQPGGPTQTNA | 9.241 | 2.491 |
| **276** | PSNSQNFSPNGQGNGPNNQDTPPRRRRRQKTRAQN EGGDTSMSGSGILAA | 8.602 | 2.851 |
| **277** | ELQNIPTSPQNFGTSQDSPQNSGGSRKRRRRGRAQSP DPAMNGGNGTNNA | 27.887 | 19.434 |
| **278** | PSQSNNGQGQSPGQGQNSNLSNPPRSRRRRKRENDF GTGMDIPTNGATAA | 1.115 | 0.732 |
| **279** | NNQLSDNGNPGSQAFDPPGGNNSRNRRRPRKRQTG STMQGIQTAESGSPA | 3.456 | 1.927 |
| **280** | TGFGDSNGQTSTIGQQPLPSQPPRKGRQRRRNRPNN SDANGSANEMSGNA | 7.735 | 2.074 |
| **281** | LQQNSPSSPDNGITSGGETGSNSRAKRRRRRFGPNMA PTGQDNPQGNNQA | 51.045 | 33.968 |
| **282** | SNSGSPSANQQETNFSPANMPINRRKRDRRQGRTQP GLNDTQGPGSGNGA | 4.495 | 2.161 |
| **283** | GSDQDNGSNTGAINETGQLPQTMRRRGRGAKRRPS NSNPSSGFPQNQNPA | 36.676 | 32.758 |
| **284** | QGNLGINSPTQQFPTGSMDESPQRKRRRAGRRAGGS PDGNNNNTNSQSPA | 1.812 | 0.437 |
| **285** | NGSNNPGTNINDNQAGQDMQSGFRRSPPRRRRKQP LPTGSNGSQAGSETA | 29.174 | 23.579 |
| **286** | NNPQQNGPQAGLNGQDNQSDIGPRRKRRRTSRGAG GNSTSNETSMFSPPA | 4.381 | 1.613 |
| **287** | SSQPSENGPFGITTGGQQPMNSPNRLRKRGRRRQTN NPGDSDNQNASAGA | 29.874 | 7.870 |

| | | | |
|---|---|---|---|
| **288** | GMDNQSSGSNNAGPDGEANSNQNRRIRKRLGRRTQ PQTGNPFGPPSQSTA | 66.032 | 46.026 |
| **289** | GNGQGNSNSGPDPSQQDMNTENLKRARSRRRGRGG NPPAFSNTQSTQPIA | 13.359 | 12.420 |
| **290** | TGSTNANNAGGNGLGIPSNGQSMNKQRRRPRRRSSD QDPQQPNPTEGFSA | 17.741 | 12.614 |
| **291** | TQPSNNSPPMQGQANGNTAGLNFRTRRRKRPRGSGS DQNSNSGPDEGIQA | 3.614 | 5.693 |
| **292** | STNDFGQINGNNSSPTNAGQDEGSKRGRRRRPRQSP MNGPGQQLNPTASA | 117.597 | 81.866 |
| **293** | SQNFNPPNLSSTPGSADQGQTGNRGRKPRRRERNM DQIQGTANPSGGNSA | 0.531 | 0.138 |
| **294** | DDIQTFQNNQSNGMGPPGLGGNQKRRRRSPSRRSAS SPTTANPEQNGNGA | 6.551 | 8.074 |
| **295** | TQAQDNQNSPGDTPSGESANSNTRGLRRRKRIRPGP GGSNNGPNFMQQSA | 21.862 | 11.567 |
| **296** | AASNTNDSQQQGIGSQNLGNGGGRNRKRRRESRPN FPQPPSNMPGDTSTA | 0.848 | 1.419 |
| **297** | NDSTQGNGPGSNQPSGESAFNQMQRRKRRPRGRNL GTQNSGITDNSAPPA | 2.778 | 0.616 |
| **298** | MPNTNGPQQEISFSNGSSNGNPTGRRRRRAKGRSGN NQAPTSQGPQDDLA | 21.380 | 3.110 |
| **299** | SGINDGTSQNLEPPMNDSQGGASRKNRRTGRRRNQ FAQNQGPPTPSSGNA | 5.451 | 2.221 |
| **300** | NNGNGQTFSGLQSGITDNNNMTQKRPRRRDGRRGP ESQGNPPAPSAQSSA | 2.757 | 2.553 |
| **301** | GSINPDNGPPSTGGGAFDNGLTQRGRRKRRNQRNQ SSPQNPSETQMNSAA | 10.007 | 7.579 |
| **302** | MNSPTQPSNDQDEAGLQNGNQSTRGRNRRKRRGPS GGPNQPSAIGNFTSA | 163.026 | 57.051 |
| **303** | DPPNNTSQNGLGEQNGQMIPTNFRRRDGTRRKRSS NSSGQPAASPGGQNA | 13.254 | 2.623 |
| **304** | SSQPNMQNNGQNLGPPNPDNNPSRIRKGRRRRGSD QTAASGFEGQTSGTA | 2.480 | 0.570 |

| | | | |
|---|---|---|---|
| **305** | LNDGTSGDQMNPPGSQPSTFAGNKQRRGNRRRRSG TQNIEGSPNANSQPA | 42.929 | 35.001 |
| **306** | QGSNSSAQNSPDSSNQPTNFPLNQRNRRRRIKRMGT GDNPPEAGQGGGTA | 10.991 | 9.071 |
| **307** | PQSDNIGQTQQLNTSNSGFAENPRKRARRGRNRPNG QSTPDPMGGSGSNA | 64.173 | 24.330 |
| **308** | TSGQNGQPFLSQNGPGTSESDPNRRRRKISRMRQNP NPASTDNNQGAGGA | 1.780 | 1.125 |
| **309** | GQDLSEGGNQQQSSPSNNSPFTDRKRRRRARGGMG GNAPNPSIQNNTPTA | 5.307 | 0.960 |
| **310** | GTSGGQNTGQSTMNSAGGLAQNNKNRRRRRNRQD NPSFSQIDEGPSPPPA | 11.873 | 10.659 |
| **311** | PSGNSPNTGSQEMLQISPDQGAGSRRRGKARRRPNF DNNQNTPTGGNQSA | 87.348 | 31.554 |
| **312** | DGFQQNISEGNGSNGQGPANDTTMRKGRRRRRQTN SPGLASNSQPNSPPA | 136.269 | 77.374 |
| **313** | LQAGFPGNSQTNQDTNTGPAPDMRKRRSGRRRSSPE SIQQGNNGGNSPNA | 12.417 | 4.498 |
| **314** | NAFSGNQGPPASQSDILQGNNGNKRPRRGDRRRTG MNTNEPSSPTGQQSA | 12.087 | 6.863 |
| **315** | QQGQGSTGNPSQLPMNTQNADTAKRNRRRIRRNFP GGPPGSSDNSSNGEA | 11.890 | 7.626 |
| **316** | SPNLNNSQSIQSSSMQFNPPDPGARRRRQKRNRNQG TGDGGTGTGNEPAA | 2.617 | 0.395 |
| **317** | QSNGPQTSNSSEIGFQNTSTSGDNKPRRNRRRRAGPN PNGMADLPQQGGA | 1.037 | 0.576 |
| **318** | ASSSNQQFSNNTQDLGIPNQPEGKRRRRRQTTRNGS PPSAMPGGNGDNGA | 8.539 | 4.873 |
| **319** | NSQNMPGGPAQGNINTEGTSGSSRDRDRRRRNKPTG NPSQQGFANPSQLA | 0.694 | 0.518 |
| **320** | ENSDGNGNAPQFQNGDPMQPQNGRGRRKSQRRRT SGGSNTTPALSSNIPA | 9.605 | 7.051 |
| **321** | TTPDANNAQGNGFLQPGQSSIGGPQGSNRNKDTRR RRRQSNGMPPENSSA | 16.724 | 30.411 |

| | | | |
|---|---|---|---|
| **322** | QMGIPNTSGPSQFGQEPNGQNANGSGTSGKNRARR RRRQNTDDSNPLPSA | 71.885 | 49.463 |
| **323** | NSGTGSQNPPGQSIAGPDNNMAGTFNGSRRDQRRQ RKRSGLPSPNQNETA | 4.311 | 2.556 |
| **324** | PNSNQGTDSPGNASNLEGNQQPQDPGSMRRKFTRR RGRASNNIPTGSGQA | 275.400 | 349.123 |
| **325** | QGGAGNQMNTSPSDSTTISPQPLGPNDPKSSRRNRRR RNQNQNEGGFGAA | 327.639 | 664.680 |
| **326** | NPQNNTFGASGGLGQSMNENNQPTSQIGKARRPRS RRRQTGSDSDNPGPA | 227.587 | 110.054 |
| **327** | NSTQIGEPASSNSTGQGGADFGMSNPNPRKRTRRQR DRPPGNNGSNLQQA | 111.354 | 79.094 |
| **328** | PNQGDSESPDGSTGNSGNNNTSPSQQPNRIGRKRRA RRTAMPFLGQGNQA | 100.893 | 94.087 |
| **329** | GGSGILGNQFNPGSATNNQDQNPQASPTNRRKRPR DRRESNPMGTGSSQA | 1.319 | 0.619 |
| **330** | GNQDDQQNSSQESINQFGPPNGSNPSASRRNRRGRT KRLTNGGPMTPAGA | 40.485 | 56.131 |
| **331** | GGGSGSDQTNAQSFGPNPEQQTSGNAIPRRMRSNKR RRDGSLTNNNQPPA | 227.283 | 161.009 |
| **332** | NMQDPQPGGQNGNLTGTSSAGPNPSFQITRRRKRER GRQSSDAGSNPNNA | 39.687 | 32.402 |
| **333** | DLNINGMNESFGPTNSGGSSQNTPQQQGRRRRNKQ RSRGPPATNDPAGSA | 69.786 | 41.415 |
| **334** | PPSGTSPSNGFGNGMQINNTQGNDNGNTRAGRRRR SKRPSQDQLSPEAQA | 1.939 | 0.654 |
| **335** | GNFSNSLQQASGPINGGTAPNDQPNNNDRTSRRRPR KRQQGTPGGSSEMA | 0.397 | 0.055 |
| **336** | QSDQGNGNPTQELNAIAPQSSSFNDSGNGPRRKTRR RRGMTSNPPGQNGA | 8167.374 | 4679.335 |
| **337** | DGANALNIGGSQNSSPGQSSEQTNPGMGKPRRRTPR RRPTFSDGQNNNQA | 176.150 | 42.024 |
| **338** | LGGANMTEGGPNTSIGDNQSNQAPSNPFRGRRKSQR RRQSNPDTSGPQNA | 122.630 | 108.895 |

| | | | |
|---|---|---|---|
| **339** | PPESSNNFNGQTTNPTISAAQGGGQNMSRKRRRRRQ PGGNDGSDNPQLSA | 86.825 | 79.913 |
| **340** | SQPSGPGSGGDTNSPPQPDGNSNGNQFEKRRNRRR NARMLTNTSQGAIQA | 55.761 | 37.267 |
| **341** | QTQGPNSSSANAGSGDTQQNSTPGNFSDIRRRRGKR RMPNQGEPGNPNLA | 73.533 | 61.964 |
| **342** | PNGSNMDDQPAGGPGNSPQQTSTEFANQRSRRKRRR NTGGQPNNLSSGIA | 49.750 | 77.367 |
| **343** | TQTPSPAGNGQFILTGEMNSPDADNQQSRRPRKRRSR GNSNQGGSPNGNA | 322.527 | 210.639 |
| **344** | SNSQTGAAPPPNFQNGGGGSTEGNGMPNRRRKQRR SRSDQNQILSNPDTA | 7.093 | 1.663 |
| **345** | TGMPPNQQNDSPNGSNFSGPGGNILGQGKRNQRQR RRRATSNSPASETDA | 19.733 | 16.108 |
| **346** | GPSNNQSSPFGDGMLISNNGTTQNPQPERTRKRRAR RGNQDGGSNASQPA | 54.745 | 72.541 |
| **347** | GSPIQGQNNSANDTNGQTSFSGSNPMAGLKRRRQRR PRSGDPTPNGEQNA | 33.377 | 37.549 |
| **348** | FNAGMNETGGSADQNSNQGGTQQSDGSLPNNKRR RRRRPPQGNSPITSPA | 1108.910 | 555.489 |
| **349** | GNSDNGQNLEGTSQPGNAPNPNPGSGTQRRQSFKR RRRSMNQSAGITPDA | 1.365 | 1.000 |
| **350** | SFPNAINGQQPQNASQDSTGPNEGNMGLRKGNRRT RRRGSNTPQSGSPDA | 6.352 | 1.696 |
| **351** | QSSMGSGNNNDGDNQPQITPGQNTLENPGARKRRR SRRGNFSQPASTGPA | 16.531 | 6.090 |
| **352** | NNGMPNQPINGQNQTGSTSQLGSFGASPRRRNKPR GRRSGPQANDESDTA | 4.504 | 2.523 |
| **353** | GPSDTTQSNQGNQPSNAILQESGNGGPGRRRSMRD KRRPTSNGPNNAQFA | 1.043 | 0.582 |
| **354** | NNQSNPSQQGAQDGNGNAPMTPGPPNNSSRFRGKR RRRGEDSILGTQSTA | 17.098 | 10.976 |
| **355** | PQQQNDDGNPSSGTTTGQNFSQMAPESNRRKPRRG RRNGNAPGNLSIGSA | 197.226 | 127.581 |

| | | | |
|---|---|---|---|
| **356** | NSSQLEPTISGNNDNGPTPAMSPNGDGNKRRSRRRAQRQPGNTGGQSQFA | 47.644 | 56.124 |
| **357** | DNQINQGNAPMPPEAGQDGGFTSSSGGSNRKRRSGRRRNNQNLSPQTPTA | 1493.647 | 317.584 |
| **358** | QNQQSNSLNNGTASNMPPGPETQGNISPGRRRRGKSRRQTDPGFGDSNAA | 2.870 | 1.526 |
| **359** | LGPQPSSQQPNSNSGIGDMNNNGFPSPTRRARGRRDRKGGNQNEASTQTA | 6.818 | 14.119 |
| **360** | GTNNEGSGNSPDGPNGITNSNMQPQPAQRPRRSKSRRRNAFGDQQSTGLA | 1848.318 | 844.283 |
| **361** | QSQTTGSGPPQGGPATGNNQGDSNMNPDRRRRRESRLKNQSPNSIGNAFA | 1.299 | 0.853 |
| **362** | PAQSNSPPGQGSNQPNTQEDGGTGNSTLRRRMRDKNRRSPSNAIGQFGNA | 1.065 | 0.503 |
| **363** | LQGSAPNNNSGMSGPPNQGDQNGDPGQNKRIRTPRRRRNSGTTASFSEQA | 67.831 | 46.433 |
| **364** | NNTPLNNIGQPTGSGSASQENPGGFPMQQRRRRRGKRDSSSANGPNTDQA | 128.774 | 36.707 |
| **365** | TNIGLNADSPNNSGSDSGQFSENGTNGSKRRGRRTRRPQQPGNPAMQPQA | 97.866 | 68.059 |
| **366** | TLPSMSGNTSAINNGPNESSNGPNPGSQRRKRRGRDPRGQTGQDFNAQQA | 70.854 | 47.715 |
| **367** | AQTDMNSSGLPSESTPGQNGQNQDNGINRRRNFRRRAKQPSGGPTPNGSA | 95.266 | 189.611 |
| **368** | NSGNGPASSSPQPPTTNNNDQGEDQGISRKRRRGQMRRGTSLNGNQPAFA | 82.836 | 57.344 |
| **369** | NDGSGFGPSPQMQQQGNQNISPPGTGSGTRKRRERNRRPSNTAALSNNDA | 163.476 | 97.261 |
| **370** | SNQPMSESSNGQNAITNGDDASNPGNQPRKRTQRRRQRGPSGTLPNFGGA | 3.273 | 2.949 |
| **371** | GQDNDPGFSGNNPQGEQPTNTSIQSNTASLSMNSAKRRRRRGRNGGPPQA | 16174.409 | 11539.709 |
| **372** | IFPNAQDTDPGSTSGTNNNNNMEGGGLSAQGSQRNQKRPRRRRPPSSGQA | 15707.192 | 8795.577 |

| | | | |
|---|---|---|---|
| **373** | NFPNMGQASENSGIDNGDPSTTNSPPGSQLGQNRQ KRRRRPARQSGGNTA | 9848.032 | 5233.482 |
| **374** | NNPNMAGANGFQGTQSNSGPGSISDESQQLPGQDR RKRRPRNRPTGTSNA | 1063.322 | 755.303 |
| **375** | MTNFETSNSSQGQPGNGGPQGNSQDSNTPLAGNRS ARKRRRQRNPDPIGA | 121820.409 | 77731.00 7 |
| **377** | SGSFTESGNSNGQSQPMNLIGAQDNGGNQPDPARRR PTRRNKRTPGNSQA | 10492.849 | 6629.336 |
| **378** | GPQNPNEGTSGGFLAQNNNAITSPPSSDQNQGTRRS NKRRRRGDPMSGQA | 2616.687 | 1836.891 |
| **379** | SGAMQPTEDNGNAPGISGGTLPQQNPNSNSNTNRR RKRPSFRRQGDGSQA | 11358.190 | 7967.531 |
| **380** | GGNSEMNPQFTADIPQNTNGNSSGSPPQNPGTLRRD NKRRRGRQGSQASA | 2501.247 | 2181.342 |
| **RBD** | ASWFTALTQHGKEDLKFPRGQGVPINTNSSPDDQIG YYRRATRRIRGGDG | 0.019 | 0.002 |
| **NTD** | MSDNGPQNQRNAPRITFGGPSDSTGSNQNGERSGAR SKQRRPQGLPNNT | 0.012 | 0.002 |
| **WT** | MSDNGPQNQRNAPRITFGGPSDSTGSNQNGERSGAR SKQRRPQGLPNNTA | 1.000 | 0.491 |
